# Autosomal suppression of sex-ratio meiotic drive influences the dynamics of X and Y chromosome coevolution

**DOI:** 10.1101/2023.09.28.559847

**Authors:** Anjali Gupta, Robert L. Unckless

## Abstract

Sex-ratio meiotic drivers are selfish genes or gene complexes that bias the transmission of sex chromosomes resulting in skewed sex ratios. Existing theoretical models have suggested the maintenance of a four-chromosome equilibrium (with driving and standard X and suppressing and susceptible Y) in a cyclic dynamic, studies of natural populations have failed to capture this pattern. Although there are several plausible explanations for this lack of cycling, interference from autosomal suppressors has not been studied using a theoretical population genetic framework even though autosomal suppressors and Y-linked suppressors coexist in natural populations of some species. In this study, we use a simulation-based approach to investigate the influence of autosomal suppressors on the cycling of sex chromosomes. Our findings demonstrate that the presence of an autosomal suppressor can hinder the invasion of a Y-linked suppressor under some parameter space, thereby impeding the cyclic dynamics, or even the invasion of Y-linked suppression. Even when a Y-linked suppressor invades, the presence of an autosomal suppressor can prevent cycling. Our study demonstrates the potential role of autosomal suppressors in preventing sex chromosome cycling and provides insights into the conditions and consequences of maintaining both Y-linked and autosomal suppressors.

## Introduction

Meiotic drivers are selfish genetic elements that manipulate transmission during gametogenesis to cheat Mendelian segregation and bias transmission in their favor. When these drivers are present on a sex chromosome in the heterogametic sex, the unequal transmission of sex chromosomes results in biased sex ratios among progeny and within populations (Jaenike, 2001; Lindholm et al., 2016). This phenomenon is known as sex-ratio meiotic drive (Jaenike, 2001; Lindholm et al., 2016). Driving X chromosomes lead to a female biased sex ratio diminishing the average fitness of the population (Bryant et al., 1982; Hamilton, 1967; James & Jaenike, 1990). As the increased transmission of driving X chromosomes relies on inhibiting the generation of functional Y-bearing sperm, there is a strong selection on the Y chromosome to counteract this effect. Suppression of X-linked drive can also evolve on autosomal loci. In a female-biased population, males will have a higher mean fitness than females (Fisher, 1930). Thus, autosomal genes that suppress X-linked drive will be more frequently inherited by male offspring. This will lead to their increased transmission in subsequent generations resulting in selection for autosomal suppressors. Hence, genomes can evolve both Y-linked and autosomal strategies to suppress such drivers. While there are several examples of sex-ratio meiotic drive systems where both autosomal and Y-linked suppressors segregate, there has not been a systematic investigation of how these suppressors might interact from a theoretical population genetic framework.

Sex-ratio meiotic drive systems can have profound impacts on genetic variation within populations. X-linked drivers are frequently linked to reduced recombination across extensive regions of the X chromosome (Charlesworth & Hartl, 1978; Dyer et al., 2007; Fuller et al., 2020; Pieper & Dyer, 2016; Prout et al., 1973). Furthermore, Y-linked suppressors usually tightly linked to the rest of the Y chromosome since there is no recombination on the Y outside of pseudo-autosomal regions. The presence of a stable equilibrium or cycling dynamics between drivers and suppressors, coupled with reduced recombination affects the patterns of genetic diversity and allele frequencies across sex chromosomes.

In many species, the genome has evolved to counteract sex-ratio meiotic drive [Table 1]. We distinguish two broad mechanistic categories of counteracting meiotic drive: resistance and suppression (Price et al., 2020). We assume resistance occurs when the target of meiotic drive (which could be nucleic acid sequence or protein) evolves to be less sensitive to the driver. In *D. melanogaster’s* autosomal Segregation Distorter (SD) system, some chromosomes are resistant because they have fewer copies of the *responder* locus making those chromosomes less sensitive to drive (Houtchens & Lyttle, 2003; Larracuente, 2014; Lyttle et al., 1986; Walker et al., 1989). On the other hand, suppressors can evolve anywhere in the genome and act by interfering with the meiotic drive machinery (the suite of proteins and nucleic acids that lead to the drive phenotype). Suppression can occur on the autosomes through loci that somehow interfere with the expression of the components of the meiotic drive system (Tao, Araripe, et al., 2007a; Tao, Masly, et al., 2007a). The Winters system of *D. simulans* is suppressed by a hairpin RNA, encoded by an autosomal locus, that interferes with the meiotic driver (Vedanayagam et al., 2021, 2023). In the instance of X-linked sex-ratio meiotic drive, the suppression mechanism can be encoded on the autosome or the Y chromosome, but resistance must occur at the Y-linked target. We will lump these two possibilities together and refer to them as suppression throughout because the mechanism does not change our modeling approach.

**Table 1:**
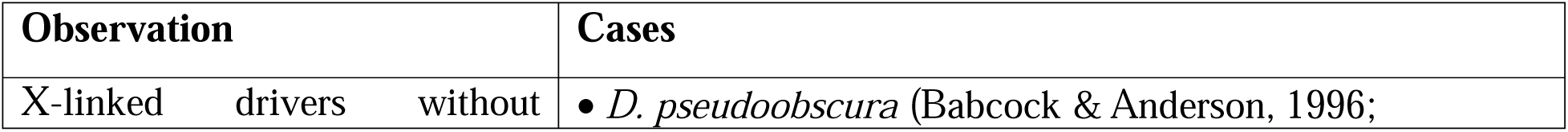
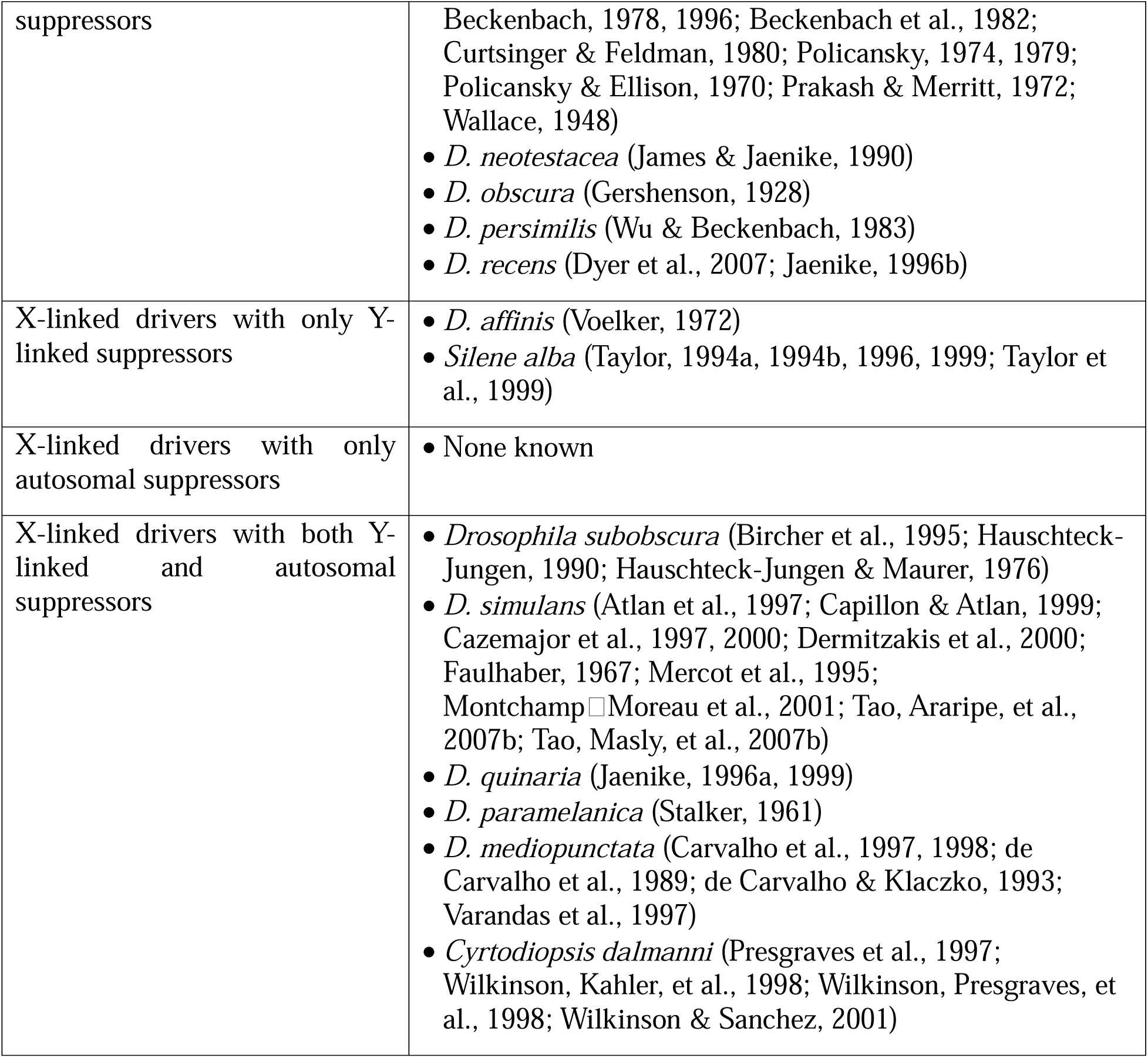
Examples of species with X-linked sex-ratio meiotic drive and its suppressors.

The selective benefits of the suppression of drive depend on the chromosome on which the suppressor resides. Any X-linked driver exhibiting strong drive and/or at high frequency will result in a female-biased population because males carrying the driver produce an excess of daughters relative to sons (Jaenike, 2001). Classic sex-ratio theory predicts that any mutation that suppresses meiotic drive, and is therefore found at higher frequencies in rare males, will have a selective advantage (Hamilton, 1967). Each offspring has one mother and one father, hence rare males are expected to produce more offspring than common females. Thus, suppressors that restore a one-to-one sex ratio are favored by selection regardless of the chromosome on which they are present as long as the sex ratio is biased. (Frank, 1991). Y-linked suppressors have an additional selective advantage since they are associated with higher mean fitness than their susceptible Y-linked counterparts (Hurst & Pomiankowski, 1991). Unlike the indirect effect on autosomes, suppression of drive confers an immediate and direct advantage to a suppressor on the Y chromosome (Burt & Trivers, 2008). Autosomes spend half of their time in females and the other half in males, therefore an autosomal suppressor is associated with a fitness advantage only half (or less if the population sex ratio is biased toward females) as frequently as a Y-linked suppressor (Rice & Holland, 1997).

There is an extensive literature within theoretical population genetics investigating the maintenance and stability of X-linked meiotic drivers (Clark, 1987; Curtsinger & Feldman, 1980; Edwards, 1961; Hall, 2004; Vaz & Carvalho, 2004; Wu, 1983). Hall, 2004 notably predicted the maintenance of a stable four-chromosome equilibrium (X^S^ – standard/non (meiotic) driving X, X^D^ – sex ratio (meiotic) driving X, Y – drive susceptible Y, Y^S^ – meiotic drive suppressing Y) when a driving X chromosome and a Y-linked suppressor are segregating in a population, and costs associated with the driving X and suppressor are small. Hall, 2004 also predicted that X^S^/X^D^ and Y/Y^S^ might undergo stable cycling under some parameter sets. Following Hall, 2004 we use the term “sex chromosome cycling” here for the cycling between a standard and driving X, and a susceptible and suppressing Y (X^S^/X^D^/Y/Y^S^). Despite these predictions from a theoretical model, field surveys focused on sex-ratio meiotic drive have failed to capture sex chromosome cycling in wild populations (Carvalho & Vaz, 1999; Dyer, 2012; Pinzone & Dyer, 2013). Hall suggested several possible explanations for the absence of cycling dynamics in natural populations, including relatively short observation timeframes, migration, and the prospect that autosomal suppressors of X-linked drivers may impede the anticipated cycling of sex chromosomes (Hall, 2004). Numerous studies have examined Y-linked suppression of drive (Clark, 1987; Hall, 2004), and autosomal suppression of drive (Vaz & Carvalho, 2004; Wu, 1983) individually but, to the best of our knowledge, only Atlan et al., 2003 considered both mechanisms of suppression of X-linked meiotic drivers in their model. However, their work does not generalize beyond *D. simulans* (where suppressor alleles show no fitness costs, maintain low quasi-equilibrium frequencies, and exhibit a prolonged period of transient polymorphism), leaving a gap in understanding the interplay of Y-linked and autosomal suppressors in broader contexts. This study is motivated by the empirical observation that many sex-ratio meiotic drive systems segregate for both Y-linked and autosomal suppression [Table 1], and the lack of theoretical treatment of co-segregating Y-linked and autosomal suppression.

We develop a mathematical model to consider the population genetics of an X-linked driver, a Y-linked suppressor, and an autosomal suppressor. Using numerical simulations, we address whether autosomal suppressors can prevent sex chromosome cycling or even the initial invasion of the Y-linked suppressor. An autosomal suppressor can effectively disrupt sex chromosome cycling, if it possesses the ability to: a) invade a population at equilibrium with a driving X and a Y-linked suppressor, and b) impede the invasion of a Y-linked suppressor in a population at equilibrium or cycling for an autosomal suppressor. If a Y-linked suppressor cannot invade the population, the cycling dynamics predicted by Hall 2004 are impossible. First, we compared populations where a driving X-chromosome segregated either with or without an autosomal suppressor to ask whether the presence of an autosomal suppressor influences the ability of a Y-linked suppressor to invade the population which we refer to as Scenario A. Next, we considered the opposite scenario: the invasion of an autosomal suppressor in populations at equilibrium for a Y-linked suppressor and the driving X in Scenario B. We specifically examined cases where the initial populations exhibit stable cycling for the driving X and Y-linked suppressor and simulated the invasion of an autosomal suppressor to see if this impeded cycling in Scenario C. Finally, we explored parameter spaces where, in the absence of autosomal suppressors, Y-linked suppressors exhibit stable cycling with the driving X, and ask whether the presence of autosomal suppressors impede such cycling in Scenario D. We close by discussing other potential factors that might explain why cycling dynamics have not been observed in natural populations.

## Methods

### The Model

We model this sex-ratio meiotic drive system by assuming a bi-allelic, tri-locus system combining elements of models developed by Wu (1983), Clark (1987) and Hall (2004). We assume an infinite population size (though we briefly relax this assumption later), discrete non-overlapping generations, diploid organisms, a single panmictic population, and all individuals have the same number of offspring. An X chromosome can be either X^S^ (standard X) or X^D^ (D: meiotic drive), a Y chromosome can be either Y (standard/susceptible Y) or Y^S^ (S: suppressor), and an autosome can be either A (standard autosome) or A^S^ (S: suppressor). A single copy of a suppressor (either A^S^ or Y^S^) is sufficient for complete suppression of the drive locus, in other words, suppression is complete and dominant. In the absence of a suppressor, an X^D^Y male produces (1/2 + d) X^D^ sperm (d represents the strength of drive) and (1/2 - d) Y sperm (0 < d ≤ 1/2). We assume viability cost for carrying a driving X or suppressors. We denote these costs as 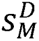 and 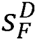 for the cost of carrying a sex-ratio driving X in males and females respectively. We denote the cost of carrying a Y-linked suppressor as *s^Y^* and an autosomal suppressor as *s^A^* which is the same regardless of sex. These costs can range between 0 to 1. Furthermore, the dominance of X^D^ on viability in females is denoted as *h^D^* and the dominance of A^S^ on viability is denoted as *h^A^*, where 0 ≤ *h* ≤ 1, however, we restrict our analysis to the three simple cases where costs are recessive (*h*=0), additive (*h=*1/2), or dominant (*h*=1). Following Wu, 1983, we denote the frequencies of the *i* genotypes for females as p_i_ and for males as q_i_ and each genotype is associated with a mean fitness (*<u>u</u>_i_* for females and *v_i_*for males, Table S1). Note that we refer to an allele to be nearly lost (and the other nearly fixed) in the simulations for our infinite population model if it consistently remained at a frequency less than 10^-16^ for >5000 generations.

**Table 2:** Variables and parameters used in the model.

| Variable / Parameter | Description |
| --- | --- |
| $X^S / X^D$ | X chromosome genotype (standard/drive) |
| $Y / Y^S$ | Y chromosome genotype (susceptible/suppressor) |
| $A / A^S$ | Autosomal genotype (susceptible/suppressor) |
| $p_i / q_i$ | Genotypic frequencies in females/ males (see Table S1) |
| $h^A / h^D$ | Dominance coefficients for viability ($A^S / X^D$ in females) |
| $s_M^D / s_F^D$ | Cost of carrying a sex-ratio driving X in males/ females |
| $s^A / s^Y$ | Cost of carrying an autosome/Y-linked suppressor |
| $d$ | Strength of drive ($0 < d \leq 1/2$) |

We tracked the genotypic frequencies across generations using recursion equations assuming an infinite population size and discrete generations in a deterministic model (see Supplementary Information). Due to the complex nature of the model, we were unable to find analytical solutions for the six-chromosome equilibrium (X^S^/X^D^, Y/Y^S^, A/A^S^). Instead, we took a deterministic simulation approach to explore the parameter space. Upon simplification to a reduced version of our model, we get the same equilibrium solutions as Wu (1983), Clark (1987) and Hall (2004) [see Data & code]. Specifically, removing autosomal suppressors and fitness cost of drive in males yields Hall’s equilibrium solution equations 7-9 for X and Y (Hall, 2004). On the other hand, removing Y-linked suppressors, assuming perfect drive (drive strength = maximum), and zero fitness cost of autosomal suppressors, yields Wu’s equilibrium solution equations 3-3’ for X and autosomes (Wu, 1983).

### Simulations

We started each simulation realization with an infinite population size initially at a 50:50 sex ratio where the driving X (X^D^) and autosomal suppressor (A^S^) were segregating at a low frequency, and Y-linked suppressor (Y^S^) was absent in the population (frequency of X^D^=0.015, Y^S^=0, A^S^=0.005). Assuming discrete generations and infinite population size in a deterministic model, we ran the simulation by calculating frequencies based on recursions (Supplementary Information, Equations 1-5) and tracked genotypic frequencies for 5000 generations. After 5000 generations, we asked whether the population reached equilibrium (we consider an equilibrium is reached if the genotypic frequencies don’t fluctuate over 500 generations (gen 4500-5000), evaluated using the variance in each of the genotypic frequencies), and if it did, we introduced a Y-linked suppressor (Y^S^) into the population at a low frequency (Y^S^=0.0005). We then ran the simulation for another 5000 generations and tracked genotypic frequencies as before. At the end of 10,000 generations, we assessed whether the Y-linked suppressor (Y^S^) invaded the population, and when it invaded, how invasion affected the population equilibrium for all three loci. We repeated this process over a range of parameter space spanning combinations of variable fitness costs for the driving X and suppressors (incrementing 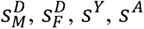 by 0.01 each time between 0 and 1), variable strength of drive (incrementing drive strength (d) by 0.01 each time within 0 to 0.5), and variable dominance (recessive, additive, or dominant).

To simplify a complicated set of simulations, we’ve divided them into four scenarios that explore the invasion likelihood, influence on equilibria and influence on cycling dynamics (Table 3).

**Table 3:**
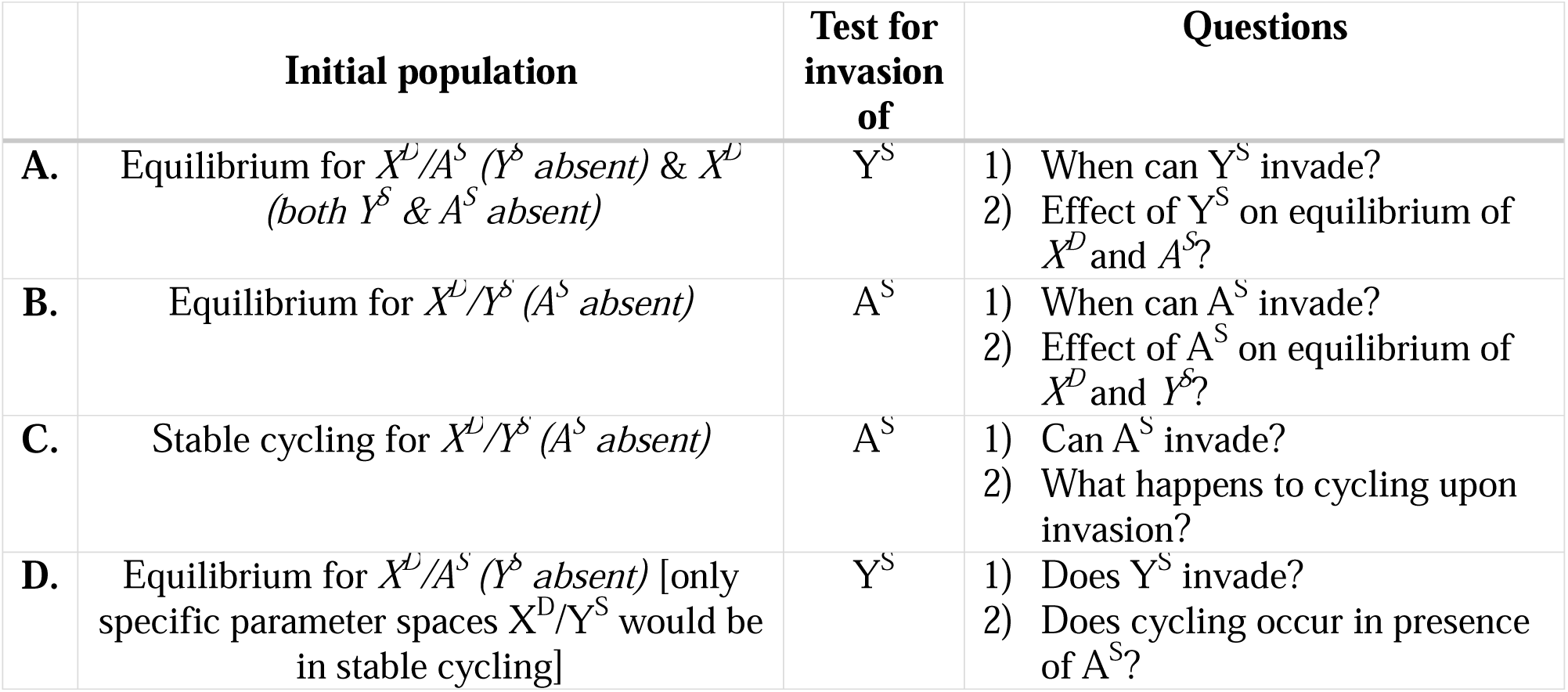
Summary of the different simulation scenarios.

|  | Initial population | Test for invasion of | Questions |
| --- | --- | --- | --- |
| **A.** | Equilibrium for $X^D/A^S$ ($Y^S$ absent) & $X^D$ (both $Y^S$ & $A^S$ absent) | $Y^S$ | 1) When can $Y^S$ invade?<br>2) Effect of $Y^S$ on equilibrium of $X^D$ and $A^S$? |
| **B.** | Equilibrium for $X^D/Y^S$ ($A^S$ absent) | $A^S$ | 1) When can $A^S$ invade?<br>2) Effect of $A^S$ on equilibrium of $X^D$ and $Y^S$? |
| **C.** | Stable cycling for $X^D/Y^S$ ($A^S$ absent) | $A^S$ | 1) Can $A^S$ invade?<br>2) What happens to cycling upon invasion? |
| **D.** | Equilibrium for $X^D/A^S$ ($Y^S$ absent) [only specific parameter spaces $X^D/Y^S$ would be in stable cycling] | $Y^S$ | 1) Does $Y^S$ invade?<br>2) Does cycling occur in presence of $A^S$? |

*Scenario A:* We first determined whether segregating autosomal suppression influences the behavior (invasion ability and equilibrium frequency) of a suppressor on the Y chromosome. To determine the effect on X and Y equilibrium frequency, we compared the simulations with the autosomal suppressor at equilibrium or absent, then introduced the suppressing Y chromosome.

*Scenario B:* To determine whether a Y-linked suppressor influences the behavior of an autosomal suppressor, we repeated the simulations but with autosomal suppressors (A^S^) initially absent (A^S^=0) and tested for the invasion of an autosomal suppressor in a population at equilibrium for the Y-linked suppressor and driving X. We again examined the parameter spaces where an autosomal suppressor could invade the population, and what happened to the population equilibrium upon invasion.

*Scenario C:* We explored cases where our initial population was in stable cycling for the driving X and Y-linked suppressor, and tested if an autosomal suppressor could invade stable cycling populations and what happened to the cycling upon invasion.

*Scenario D:* We directly addressed cases of cycling between X and Y genotypes described in Hall, 2004. In these simulations, we began with equilibrium for the driving X and autosomal suppressor and determined whether the Y-linked suppressor could invade and whether cycling still occurred.

The simulations were written and run in R version 4.2.2 (R Core Team, 2013). Data was analyzed and plotted in R using the following packages: ggplot2 (Wickham, 2011), ggpubr (Kassambara & Kassambara, 2020), and wesanderson (Ram & Wickham, 2018). All data and code is available on Github: https://github.com/anjaligupta1210/AutosomalSuppressionOfMeioticDriveCanPreventSexChromosomeCycling.git.

## Results

### Scenario A: Autosomal suppressors can prevent the invasion of Y-linked suppressors

We first considered whether a Y-linked suppressor could invade a population at equilibrium for both X-linked driver and autosomal suppressors. We compared the dynamics for the invasion of a Y-linked suppressor between populations where an autosomal suppressor was initially absent or present and at stable equilibrium to determine whether the presence of the autosomal suppressor influenced invasion of the Y-linked suppressor. When the fitness costs of the driving X and autosomal suppressor were recessive, and there was no fitness cost of carrying a driving X in males, a Y-linked suppressor could not invade a population where an autosomal suppressor was at equilibrium unless the cost of Y-linked suppression was relatively low [Figure 1]. The yellow space in Figure 1 are the sets of parameters where autosomal suppressors at equilibrium inhibit the invasion of Y-linked suppressors. We were able to obtain the limiting conditions for invasion of a Y-linked suppressor. From our simulations, we found that a Y-linked suppressor was successful in invasion when *s^Y^* < 2 × *d* and 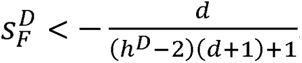. This was obtained following solutions from the reduced version of our model, and it is consistent with the results presented in Hall (2004) [see Supplementary information for more information].

**Figure 1:**
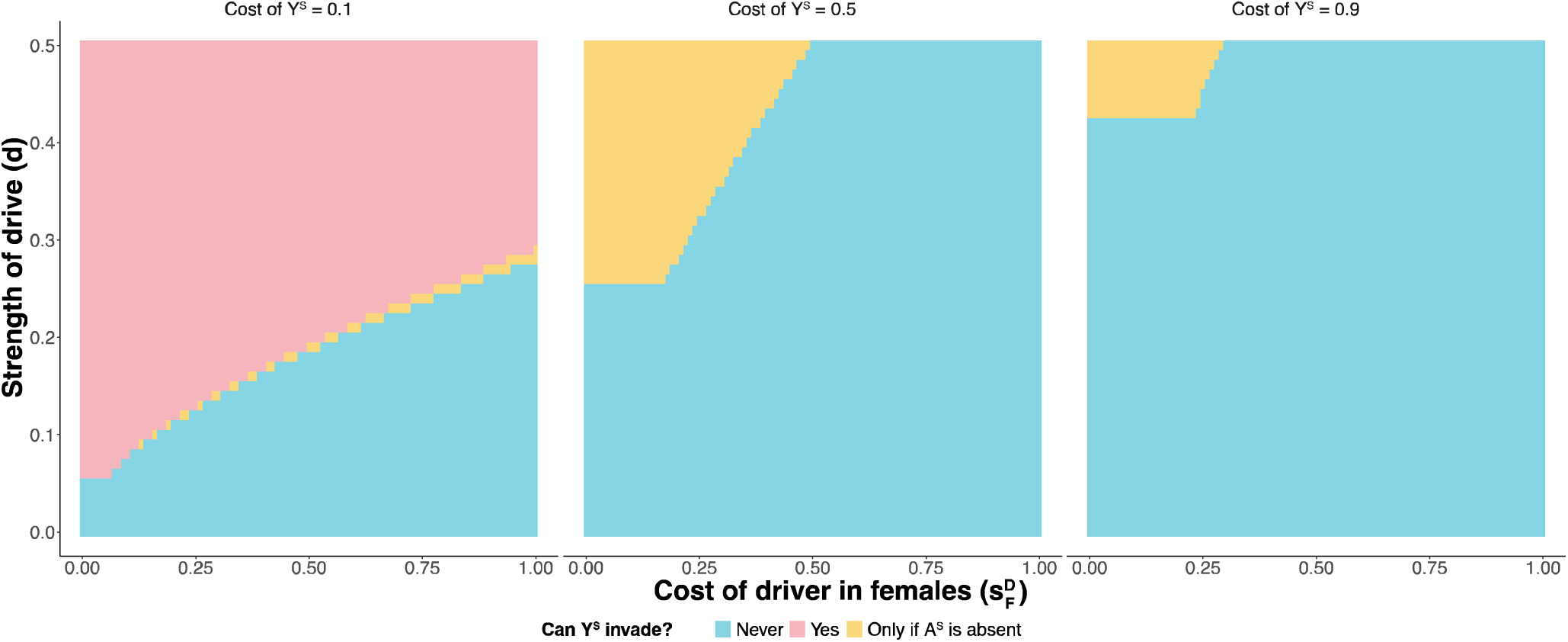
Y-linked suppressors invade a population at equilibrium for driving X and an autosomal suppressor. [*h^A^*=0, *h^D^*=0 (recessive costs), 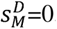] Parameter space representing whether Y-linked suppressor can invade into the population.

It is reasonable to assume that the ability of any suppressor to invade a population would be dependent on the equilibrium of frequency of the driving X. We therefore looked at whether the ability of a Y-linked suppressor to invade is associated with the equilibrium frequency of driving X (X^D^) in males before introduction of the Y-linked suppressor. Overall, when the driving X is at a high frequency, the Y linked suppressor is likely to invade, but other factors including the cost of suppression, the cost of the driving X and the strength of drive are also important. While the presence of a driving X was, of course, necessary for a Y-linked suppressor to increase in frequency, the presence of an autosomal suppressor in the population prevented a Y-linked suppressor associated with a high fitness cost to increase in frequency even when the driving X frequency was close to fixation [Supplementary Figure S1].

Since the equilibrium frequency of the driving X alone did not adequately explain the pattern of invasion of a Y-linked suppressor, we visualized the invasion ability against both the equilibrium frequency of the driving X and of the autosomal suppressor. We found that Y-linked suppressor could invade when the equilibrium frequency of driving X was high, and the equilibrium frequency of autosomal suppressor was low [Supplementary Figure S2]. When there is no cost of autosomal suppressor in the population, the autosomal suppressor rises in frequency and keeps the driving X in at lower frequency, inhibiting the invasion of a Y-linked suppressor. However, the autosomal suppressor never came close to fixation, instead it stayed at an equilibrium once the sex-ratio was restored to 50% females. This is expected since an autosomal suppressor carries no fitness advantage when population sex ratios are 50:50.

When the cost of the driving X was dominant, the ability of invasion of a Y-linked suppressor was low. The fitness cost of the driving X prevents it from reaching a high frequency in the population [Supplementary Figure S3], and with a low frequency of drive, the benefit of a Y-linked suppressor is also low. When the costs of autosomal suppression were additive or dominant, a Y-linked suppressor with small fitness costs could invade the population because of its fitness advantage over the autosomal suppressor [Supplementary Figure S4].

For cases where a Y-linked suppressor (small fitness cost of Y^S^) could invade a population with an autosomal suppressor at equilibrium, we explored how this affects the equilibrium frequencies of both driving X and autosomal suppression. We looked at the relative reduction in the equilibrium frequencies of X^D^ and A^S^ in males (defined as 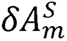 and 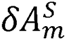 respectively) upon invasion of a Y-linked suppressor (Y^S^). The relative reduction in the equilibrium frequency of X^D^ in males 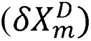 can be defined as the difference in the equilibrium frequency of the driving X before and after invasion 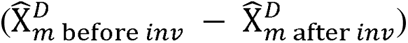 divided by the equilibrium frequency of the driving X before invasion 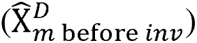. The relative reduction in the equilibrium frequency of A^S^ in males 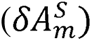 can be defined as the difference in the equilibrium frequency of the autosomal suppressor before and after invasion 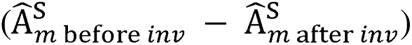 divided by the equilibrium frequency of the autosomal suppressor before invasion 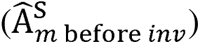

In cases of a driving X with no cost, Y^S^ did not reduce the X^D^ frequency in the population. For all other cases, invasion of the Y-linked suppressor (Y^S^) led to a decline in the equilibrium frequency of the driving X (X^D^) [Figure 2A]. The decline in the equilibrium frequency of the driving X was not monotonic for two reasons: (i) a low cost driving X could rise to a high frequency regardless of the presence of an autosomal or Y-linked suppressor, therefore the relative reduction in the equilibrium frequency of driving X was nearly zero, (ii) the decreasing gradient of the equilibrium frequency of driving X along the cost of driver in females showed some inconsistency because there were some parameter spaces where the Y-linked suppressor showed cycling after invasion [Supplementary Figure S5]. When a low-cost Y-linked suppressor invaded the population, the equilibrium frequency of a costly autosomal suppressor declined to nearly zero [Figure 2B]. We also looked at whether the dominance of costs of the autosomal suppressor has an influence on the relative reduction in the equilibrium frequency of the driving X and the autosomal suppressor [Supplementary Figure S6]. As expected, a Y-linked suppressor with a small fitness cost was able to effectively replace a dominant autosomal suppressor.

**Figure 2:**
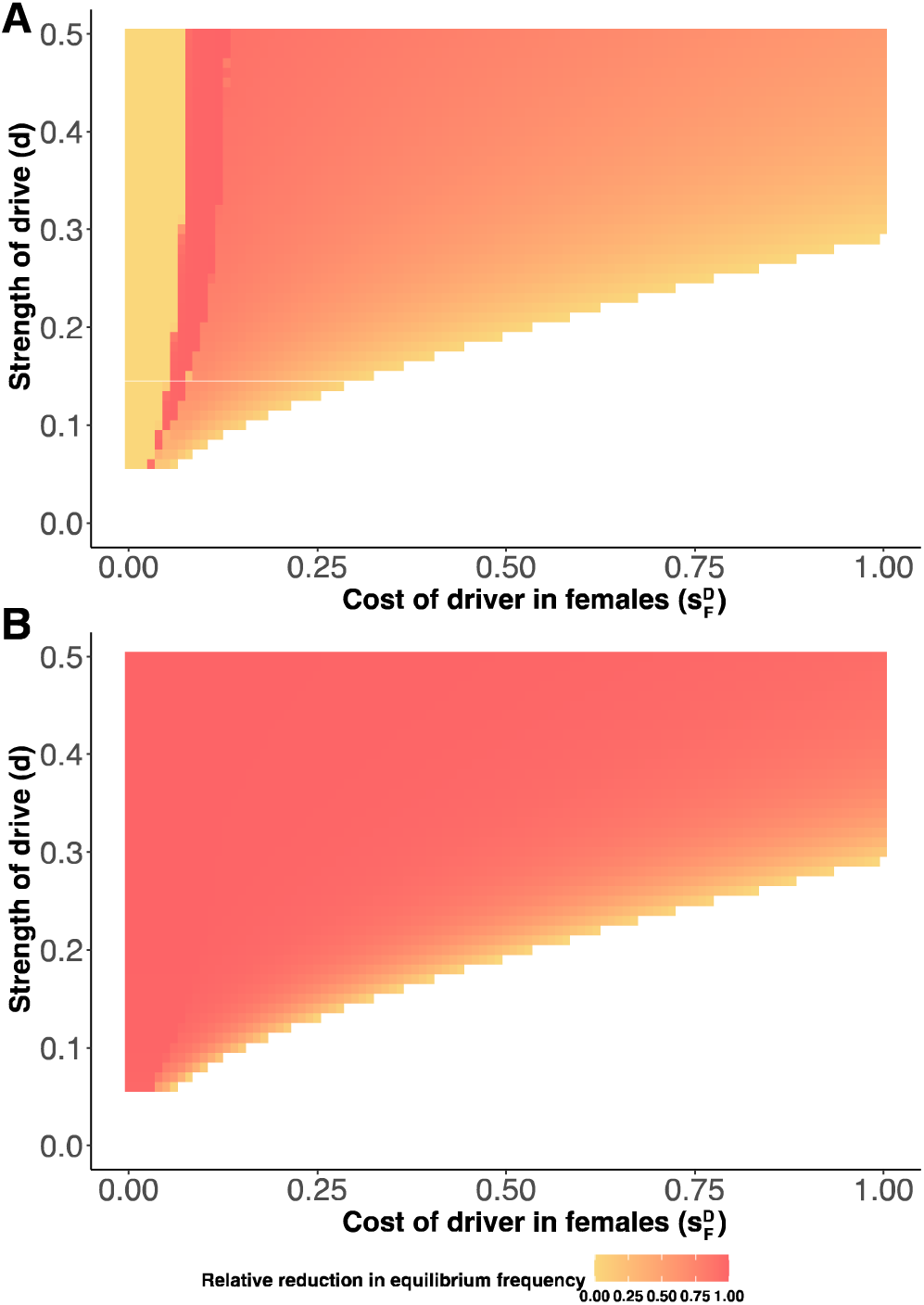
Relative reduction in equilibrium frequencies upon invasion of a Y-linked suppressor. [*h^A^*=0, *h^D^*=0 (recessive costs), 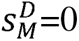, *s^A^*=0.5, *s^Y^*=0.1] **A**. Relative reduction in equilibrium frequency of X^D^ in males (see Supplementary Figure S5 for more information). **B**. Relative reduction in equilibrium frequency of A^S^ in males. The white space represents the space where Y^S^ cannot invade into the population.

### Scenario B: An autosomal suppressor can invade a population at equilibrium for the driving X and Y-linked suppressor when the sex-ratio is female-biased

In general, autosomal suppressors of sex-ratio meiotic drive are selected for because of Fisherian selection for equal sex-ratios, not because they gain a direct benefit from suppressing drive (Crow, driving X and the Y-linked suppressor were both at equilibrium *only if* the population sex-ratio was still female-biased. To test this, we ran another set of simulations starting with an initial population (50:50 sex-ratio) with frequencies X^D^=0.015, Y^S^=0.005, A^S^=0 and tracked the genotypic frequencies for 5000 generations and asked if the system reached equilibrium. When the population was at equilibrium for the driving X and Y-linked suppressor, we introduced an autosomal suppressor into the population at a very low frequency (A^S^=0.0005) and ran the model for another 5000 generations to see if the autosomal suppressor could invade the population in the presence of a Y-linked suppressor. We confirmed that an autosomal suppressor could invade the population in the presence of a Y-linked suppressor for some of the parameter space [Figure 3]. The invasion pattern of an autosomal suppressor can be broadly divided into three regions, where it invades into a population in region II, and doesn’t invade in region I and III [Figure 3, panel 4]. The invasion potential of an autosomal suppressor depended on a combination of factors – (i) the sex ratio of the population, (ii) fitness advantage of autosomal suppressor over the Y-linked suppressor, (iii) the equilibrium frequency of the driving X and the Y-linked suppressor. Regions I and II had female-biased sex ratios in the populations. All populations in region I [Figure 3] were at equilibrium for a Y-linked suppressor at a frequency close to fixation, and this Y-linked suppressor had a lower fitness cost compared to the autosomal suppressor [Supplementary Figure S8]. An autosomal suppressor could invade into the population only in region II [Figure 3, Supplementary Figure S7]. This is because the invasion potential of an autosomal suppressor is also influenced by the equilibrium frequencies of the driving X and the Y-linked suppressor coupled with the fitness advantage of the Y-linked suppressor over the autosomal suppressor [Supplementary Figure S8]. Populations in region III [Figure 3] had a driving X and a Y-linked suppressor at equilibrium at very low frequencies and balanced sex-ratios in the population, which altogether prevented the invasion of an autosomal suppressor [Supplementary Figure S7, S8]. This was not surprising as autosomal suppressors are selected for in a population only due to Fisherian selection for balanced sex-ratios. Thus, in a population with unbalanced sex-ratios, the autosomal suppressor couldn’t invade the population when (i) the frequency of driver was very low, and (ii) the Y-linked suppressor was at a high frequency. An autosomal suppressor was only able to invade a population in region II since the sex-ratios were unbalanced, and the equilibrium frequencies of the driving X and the Y-linked suppressor were not very low and very high respectively.

**Figure 3:**
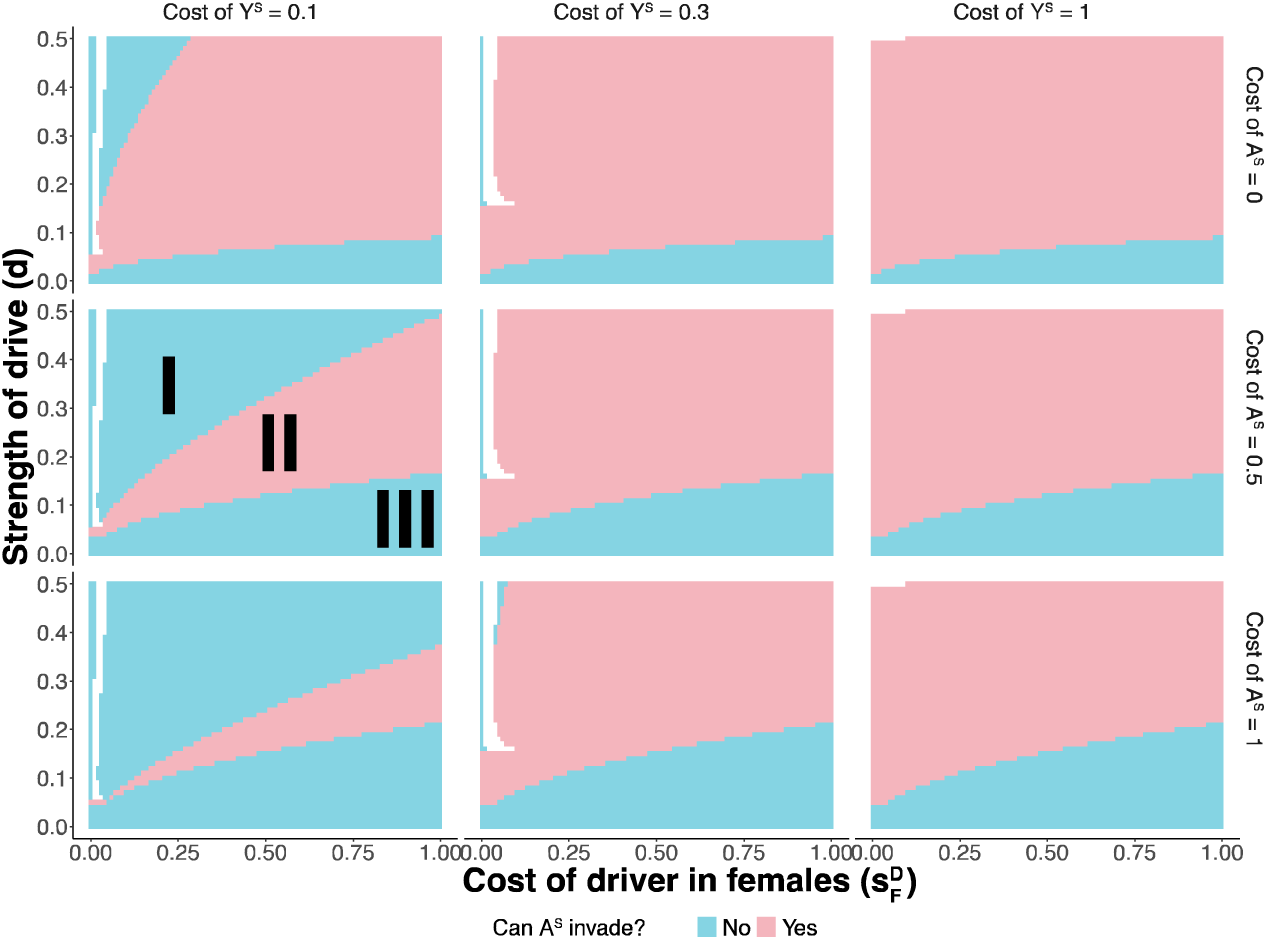
Can an autosomal suppressor invade into a population at equilibrium for a Y-linked suppressor and driving X? [*h^A^*=0, *h^D^*=0 (recessive costs), 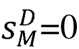, *s^A^*=0 (no 1991). We therefore cost), 0.5, 1, *s^Y^*=0.1, 0.3, 1]. (Note that the white boxes are the suspected that autosomal suppressors would be able to invade systems where the spaces where cycling occurs, and populations do not reach equilibrium as described by Hall 2004 and we deal with these parameter subsets separately in Scenario C). Regions I, II and III in panel five have been labelled by division of the parameter space based on whether autosomal suppressor invades or not. For all the unlabeled panels, the region where autosomal suppressor invades would be region II and region I and III would be to the left (or above) and right (or below) of region II respectively.

Next, we looked at the relative reduction in the equilibrium frequencies of the Y-linked suppressor and driving X (defined as 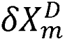 and 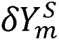 respectively) upon invasion of the autosomal suppressor. The relative reduction in the equilibrium frequency of X^D^ in males 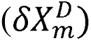 can be defined as the difference in the equilibrium frequency of the driving X before and after invasion 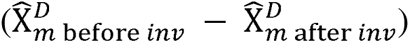 divided by the equilibrium frequency of the driving X before invasion 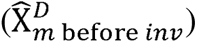. The relative reduction in the equilibrium frequency of Y^S^ in males 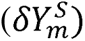 can be defined as the difference in the equilibrium frequency of the Y-linked suppressor before and after invasion 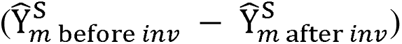 divided by the equilibrium frequency of the Y-linked suppressor before invasion 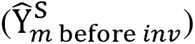.

Unlike the Y-linked suppressor in scenario A, when an autosomal suppressor invaded the population, it did not eliminate the Y-linked suppressor completely from the population, but only caused a small decline in its equilibrium frequency to restore a balanced sex-ratio. This result is important as it emphasizes the fact that autosomal suppressors are selected only selection towards balanced sex-ratios in a population. The patterns for the relative reduction in the equilibrium frequencies of the driving X showed a significant decline only when the autosomal suppressor had a zero cost and Y-linked suppression was quite costly. For all other cases, the decline in driving X was small such that the sex-ratio restored to 50-50 [Supplementary Figure S9]. In certain instances, the frequency of the driving X increased during the autosomal suppressor invasion. This occurred when the X-linked driver was initially at a low frequency in equilibrium with the Y-linked suppressor. The skewed sex ratio enabled the autosomal suppressor to invade despite the presence of the Y-linked suppressor, resulting in a decrease in the frequency of the Y-linked suppressor in the population. This, in turn, allowed a small increase in the driving X frequency until the point where the sex-ratio was balanced. These results stress the point that autosomal suppressors evolve only due to selection towards a balanced sex-ratio while a Y-linked suppressor are selected for both population level (rarer sex has higher fitness) and chromosome level (Y chromosomes with suppressors survive, those without don’t) reasons.

### Scenario C: Autosomal suppressors can invade populations undergoing stable cycles for X-linked driver and Y-linked suppressor and this impedes cycling

In simulations where we aimed to test the invasion of an autosomal suppressor in populations at equilibrium for X-linked driver and Y-linked suppressor, we had a small subset of parameter space where the driving X and Y-linked suppressor exhibited a cyclic dynamic in the absence of an autosomal suppressor as predicted by Hall, 2004. We explored this subset of parameters on a case-by-case basis, and looked at how the introduction of an autosomal suppressor affected the cycling dynamics in these scenarios. Given the cycling nature of the system, population sex ratios will also cycle, facilitating the invasion of the autosomal suppressor. In our simulations, the autosomal suppressor always invaded these cycling populations but was subsequently nearly lost in certain cases after invasion when it had a fitness disadvantage relative to the Y-linked suppressor. We observed some instances of alternative cycling of the autosomal and Y-linked suppressor with the driving X in populations when the fitness of autosomal and Y-linked suppressor was comparable [e.g.: Supplementary Figure S10-C, D, E]. Often these cycles were extreme and resulted in long spans of time when the driving X or Y-linked suppressor were at very low frequencies. In these cases, the autosomal suppressor was not really tested against the presence of the Y-linked suppressor until the Y-linked suppressor’s frequency began to increase. During the time of introduction of autosomal suppressor into the population, the Y-linked suppressor was present in the population at a very low frequency due to initial cycling, therefore the autosomal suppressor could easily invade the population. When the frequency of the Y-linked suppressor began to increase again due to cycling, the autosomal suppressor that invaded the population either dampened the cycling or impeded it by itself starting to cycle in the population.

When the Y-linked suppressor had a fitness advantage over the autosomal suppressor, eventually the autosomal suppressor was nearly lost from the population even after the invasion, and the cycling of driving X and Y-linked suppressor was impeded only for a short timespan [e.g.: Supplementary Figure S10-A, B]. But for cases where Y-linked suppression was costly and an autosomal suppressor had zero cost, the autosomal suppressor eliminated the Y-linked suppressor from the population, thus interrupting the cycling [e.g.: Figure S10-F]. In all these cases when an autosomal suppressor invaded a cycling population, the invasion impeded cycling of Y-linked suppressor and the driving X for different timespans depending upon the fitness of both suppressors [Supplementary Figure S10]. These results suggest that the introduction of a low-cost autosomal suppressor may inhibit cycling of X and Y chromosomes.

### Scenario D: Autosomal suppressors can prevent the stable cycling of Y-linked suppressors and driving X

We examined specific examples presented in Hall 2004 where a driving X and a Y-linked suppressor stably cycle. These regions of stable cycling were restricted to a subset of the parameter space where the costs of drive and suppression is small, and suppressors show complete (or high levels of) suppression. First, we tested whether a Y-linked suppressor could invade populations at equilibrium for an autosomal suppressor and the driving X in this subset of parameter space where otherwise a driving X and a Y-linked suppressor would be under stable cycling. Next, we examined whether X/Y cycling was able to become established upon invasion of the Y-linked suppressor in these populations which were initially at equilibrium for an autosomal suppressor. For most of the parameter space, the Y-linked suppressor could not invade the population and hence, cycling did not occur. There were some cases where the Y-linked suppressor could invade the population but cycling still didn’t occur in the presence of the autosomal suppressor. The parameter space where the cycling still did occur in the presence of an autosomal suppressor shrank to a very small fraction [Figure 4]. Therefore, the presence of autosomal suppressors winnowed Hall’s predicted parameter space for stable cycling considerably.

**Figure 4:**
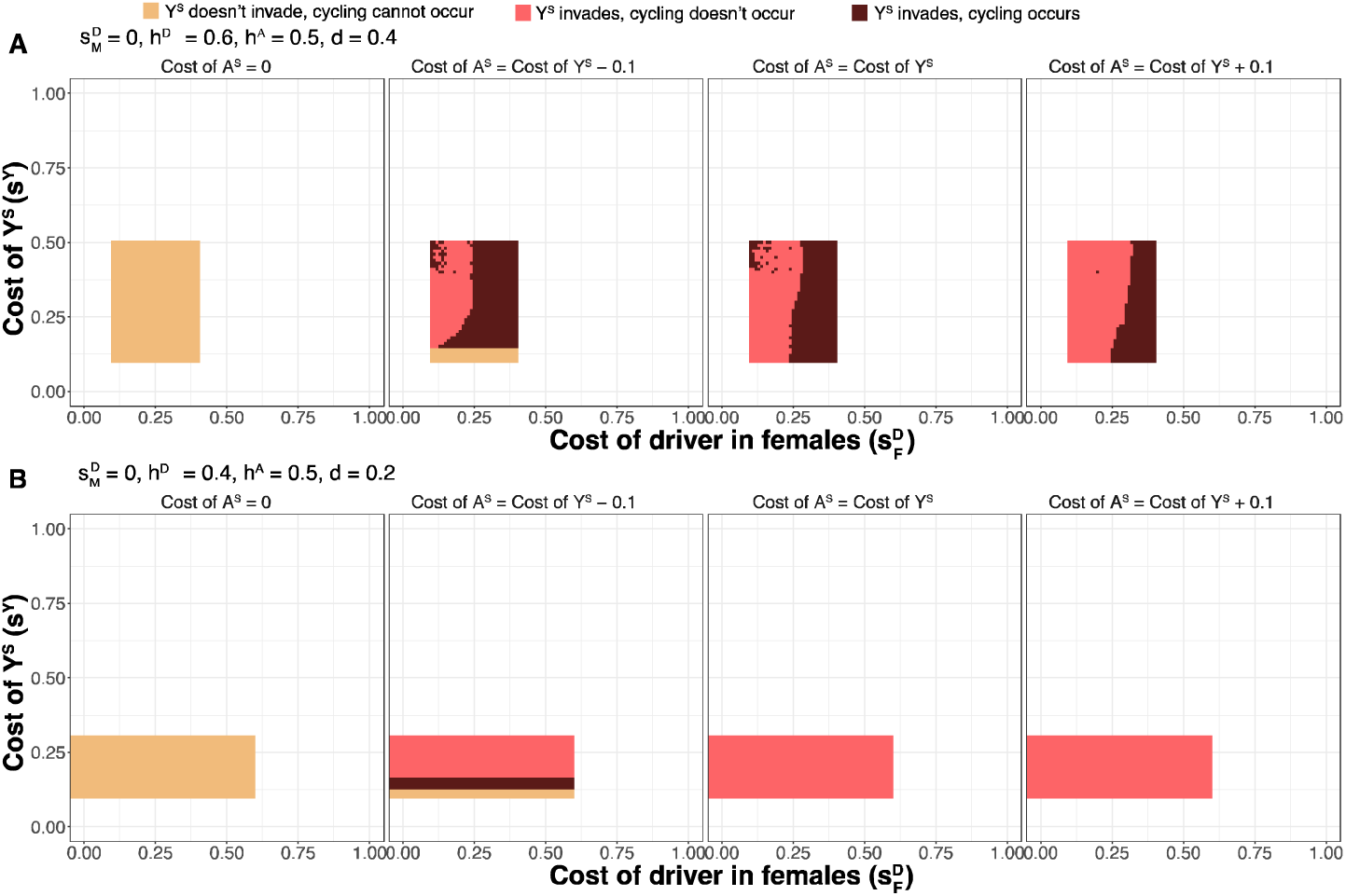
Can cycling persist in presence of an autosomal suppressor? Examples of regions of parameter spaces of stable cycling from Hall 2004, where we tested whether a Y-linked suppressor could invade populations at equilibrium for an autosomal suppressor and the driving X. The cantaloupe color boxes represent the cases where a Y-linked suppressor couldn’t invade the population and thus, cycling couldn’t occur. The salmon color boxes represent the cases where a Y-linked suppressor invades the population, but cycling still couldn’t occur due to the presence of an autosomal suppressor. The dark brown color boxes represent the cases where the cycling could persist in the presence of an autosomal suppressor. In the absence of autosomal suppressors, cycling is restricted to the shaded region.

## Discussion

Theoretical population genetic models suggest that X-linked meiotic drivers and Y-linked suppressors can exhibit stable cycling (Hall, 2004), but cycling has not been reported in any studies of natural populations. Hall, 2004 modeled how migration between populations might prevent this cycling and suggested that evolution at autosomal loci may also prevent it. Using a simulations-based approach, we show that segregating autosomal suppressors interfere with this sex chromosome cycling of X-linked meiotic drivers and Y-linked suppressors.

*D. simulans* provides an excellent case to explore how our model can illuminate the dynamics of X-linked meiotic drivers and its suppressors. *D. simulans* is one of the few well-studied meiotic drive systems harboring three X-linked meiotic drivers (named Paris, Winters and Durham) and multiple autosomal and Y-linked suppressors (Atlan et al., 1997, 2003; Cazemajor et al., 1997; Montchamp Moreau et al., 2001; Tao, Masly, et al., 2007b, 2007a). Strong Y-linked suppressors co-exist with autosomal suppressors of the Paris system in East African populations (Madagascar, Réunion and Kenya) of *D. simulans* (Atlan et al., 2003). Atlan et al., 2003 suggested that in the Paris sex-ratio meiotic drive system of *D. simulans,* autosomal suppression evolved before Y-linked suppression. This is supported by empirical observations: Y chromosomes from East African populations show complete drive suppression, while in West Indies populations, drive suppression is predominantly mediated by autosomes (Montchamp-Moreau et al., 2001).

The evolution of suppressors for the Paris sex-ratio meiotic drive system of *D. simulans* can be modelled like our scenario A or D. In such a situation, our simulations suggest that a Y-linked suppressor could invade the population under specific conditions but would rarely exhibit stable cycling unless the cost of Y-linked suppression is minimal. This aligns with Hall 2004’s predictions for stable cycling with low fitness costs of X^D^ and Y^S^. The autosomal suppressor acts as a barrier against the invasion of Y-linked suppressor. For X-linked meiotic drive systems where complete suppression of drive evolved on an autosomal locus faster than a Y-linked locus, stable cycling can rarely occur. This also raises the question whether selection for balanced sex ratios on autosomal loci may be stronger than on the Y chromosome.

Our simulations also suggest that when an autosomal suppressor has additive or dominant fitness costs, the invasion of a Y-linked suppressor could significantly reduce the autosomal suppressor’s frequency, potentially eliminating it from the population. Thus, it can be speculated that the autosomal suppressors present in *D. simulans* have low and/or recessive fitness costs. Although fitness measurements have not been made in *D. simulans*, autosomal suppressors are assumed to be deleterious in this species to account for the fact that they have not been fixed (Jutier et al., 2004). Additionally, it is important to note that other meiotic drive systems, like the Durham or Winters sex-ratio systems, might interfere with the Paris system in *D. simulans*.

For a parallel scenario, let us consider a meiotic drive system where Y-linked suppressors evolved before autosomal suppressors. Although no known empirical example of such a system exists, this doesn’t mean they don’t occur, as most meiotic drive systems remain underexplored. Autosomal suppressors could invade populations with skewed sex ratios, even if Y-linked suppressors are present. If a population is female-biased and the fitness costs of suppression are outweighed by the advantage of being in the rarer sex, Fisherian sex-ratio selection would favor the invasion of autosomal suppressors. This is indicative of our scenario B and C [Figure 3]. In such cases, it should be noted that cycling only occurs in certain restricted parameter spaces even when the autosomal suppressor is initially absent in the population. Like our scenario C, an autosomal suppressor would be able to invade in a population for most of this restricted parameter space and either impede or dampen the stable cycling of driving X and the Y-linked suppressor. Since the invasion of the autosomal suppressor is strongly dependent on the sex-ratio of the population, if a strongly selected Y-linked suppressor can completely suppress drive and nearly reach fixation in a population to restore a balanced sex-ratio, the autosomal suppressor would not be able to invade, and stable cycling might persist.

Our model suggests stable cycling of driving X chromosomes and Y-linked suppressors is a rare phenomenon. The dynamics of stable cycling are highly sensitive to specific parameters, and the presence of autosomal suppressors may significantly constrain the occurrence of stable cycling. Our findings demonstrate that autosomal suppressors can disrupt cycling in populations for varying durations depending on the suppressor fitness. Thus, even if the population eventually regains the sex chromosome cycling dynamics, these occasional perturbations may be enough to throw off cycling long enough to preclude any observation of stable cycling. We noted that some of the predicted parameter space allowing for X/Y cycling in our simulations led to frequencies that were very close to the boundaries (zero or one). We therefore considered a scenario of finite population sizes under stable cycling conditions and found that cycling disappeared due to loss of one allele in most cases with moderate (1,000,000) population sizes. Note that these simulations were slightly different to incorporate the finite population size [Supplementary Figure S11].

Interference between different modes of suppression of sex-ratio meiotic drive, specifically Y-linked and autosomal suppressors, plays a crucial role in the fates of the driver and both suppressors. Low-cost suppressors segregating on autosomes can impede the invasion of Y-linked suppressors and an invading low cost autosomal suppressor can reduce or even replace a more costly Y-linked suppressor previously at equilibrium. The opposite is also true: Y-linked suppressors can impede the invasion of or reduce the frequency of autosomal suppressors. Prior work (Hall, 2004) predicted parameter space where a driving X and suppressing Y should cycle but argued that this is rarely observed in nature. Several scenarios such as finite population sizes and migration make the occurrence of stable cycling unlikely, but here we show that the presence of autosomal suppressors shrinks the parameter space for cycling to almost nothing. Stable sex chromosome cycling due to meiotic drive may therefore be an elegant theoretical result but is burdened by so many assumptions that it is rare and difficult to empirically detect in natural populations.

## Acknowledgements

We thank members of the Unckless lab and two anonymous reviewers for helpful comments and suggestions on early drafts. We would like to thank Dr. Maria Orive for motivating AG and teaching a course on population genetics that equipped her with the necessary knowledge to pursue this research. This work was supported by the National Science Foundation Career Award #2047052 to RLU. Computational work was performed at the HPC facilities operated by the Center for Research Computing at the University of Kansas supported in part through the National Science Foundation MRI Award #2117449.

