## Supplementary Information for "Autosomal suppression of sex-ratio meiotic drive influences the dynamics of X and Y chromosome coevolution"

**The Model:**

We model this meiotic drive system through a biallelic tri-locus framework for diploid organisms. We assume an infinite population size and a single panmictic population. All individuals produce equal number of offspring across all generations. We tracked the genotypic frequencies across generations using recursion equations assuming discrete generations in a deterministic model. Upon simplification, our model reduces to models of Wu (1983) (when a Y-linked suppressor is absent), Clark (1987) and Hall (2004) (when an autosomal suppressor is absent).

**Table S1:** Genotypic frequencies, relative fitness, and the proportion of X-bearing gametes produced by them as used in the model. The frequencies/fitness for females are denoted by p_i_ and u_i_ and males as q_i_ and v_i_ (Assume normal mendelian segregation for gametogenesis in females, driving X (X^D^) disrupts mendelian segregation only in males).

| **Genotype** | **Frequency** | **Fitness** | **Proportion of X-bearing gametes** |
| --- | --- | --- | --- |
| X^S^X^S^AA | p_1_ | u_1_ = 1 | 1 |
| X^S^X^S^AA^S^ | p_2_ | u_2_ = 1 - $h^{A}s^{A}$ | 1 |
| X^S^X^S^A^S^A^S^ | p_3_ | u_3_ = 1 – $s^{A}$ | 1 |
| X^S^X^D^AA | p_4_ | u_4_ = 1 - $h^{D}s_{F}^{D}$ | 1 |
| X^S^X^D^AA^S^ | p_5_ | u_5_ = (1 - $h^{A}s^{A}$)(1 - $h^{D}s_{F}^{D}$) | 1 |
| X^S^X^D^A^S^A^S^ | p_6_ | u_6_ = (1 – $s^{A}$)(1 - $h^{D}s_{F}^{D}$) | 1 |
| X^D^X^D^AA | p_7_ | u_7_ = 1 – $s_{F}^{D}$ | 1 |
| X^D^X^D^AA^S^ | p_8_ | u_8_ = (1 - $h^{A}s^{A}$)(1 – $s_{F}^{D}$) | 1 |
| X^D^X^D^A^S^A^S^ | p_9_ | u_9_ = (1 – $s^{A}$)(1 – $s_{F}^{D}$) | 1 |
| X^S^YAA | q_1_ | v_1_ = 1 | ½ |
| X^S^YAA^S^ | q_2_ | v_2_ = 1 - $h^{A}s^{A}$ | ½ |
| X^S^YA^S^A^S^ | q_3_ | v_3_ = 1 – $s^{A}$ | ½ |
| X^D^YAA | q_4_ | v_4_ = 1 – $s_{M}^{D}$ | ½ + d |
| X^D^YAA^S^ | q_5_ | v_5_ = (1 - $h^{A}s^{A}$)(1 – $s_{M}^{D}$) | ½ |
| X^D^YA^S^A^S^ | q_6_ | v_6_ = (1 – $s^{A}$)(1 – $s_{M}^{D}$) | ½ |
| X^S^Y^S^AA | q_7_ | v_7_ = 1 – $s^{Y}$ | ½ |
| X^S^Y^S^AA^S^ | q_8_ | v_8_ = (1 - $h^{A}s^{A}$)(1 – $s^{Y}$) | ½ |
| X^S^Y^S^A^S^A^S^ | q_9_ | v_9_ = (1 – $s^{A}$)(1 – $s^{Y}$) | ½ |
| X^D^Y^S^AA | q_10_ | v_10_ = (1 – $s_{M}^{D}$)(1 – $s^{Y}$) | ½ |
| X^D^Y^S^AA^S^ | q_11_ | v_11_ = (1 - $h^{A}s^{A}$)(1 – $s_{M}^{D}$) (1 – $s^{Y}$) | ½ |
| X^D^Y^S^A^S^A^S^ | q_12_ | v_12_ = (1 – $s^{A}$)(1 – $s_{M}^{D}$) (1 – $s^{Y}$) | ½ |

The genotypic frequencies in t+1 generation, were calculated using the following recursion equations following Wu (1983):

| *T* q_12_ _[t+1]_ = $(\frac{p_{5}}{4}+\frac{p_{6}}{2}+\frac{p_{8}}{2}+p_{9})(\frac{q_{11}}{4}+\frac{q_{12}}{2}+\frac{q_{8}}{4}+\frac{q_{9}}{2})v_{12}$ | [1] |
| --- | --- |
| *T* q_11_ _[t+1]_ = $((\frac{p_{5}}{4}+\frac{p_{6}}{2}+\frac{p_{8}}{2}+p_{9})(\frac{q_{10}}{2}+\frac{q_{11}}{4}+\frac{q_{7}}{2}+\frac{q_{8}}{4})+(\frac{p_{4}}{2}+\frac{p_{5}}{4}+p_{7}+\frac{p_{8}}{2})(\frac{q_{11}}{4}+\frac{q_{12}}{2}+\frac{q_{8}}{4}+\frac{q_{9}}{2}))v_{11}$ | [2] |
| *T* q_10_ _[t+1]_ = $(\frac{p_{4}}{2}+\frac{p_{5}}{4}+p_{7}+\frac{p_{8}}{2})(\frac{q_{10}}{2}+\frac{q_{11}}{4}+\frac{q_{7}}{2}+\frac{q_{8}}{4})v_{10}$ | [3] |
| *T* q_9_ _[t+1]_ = $(\frac{p_{2}}{2}+p_{3}+\frac{p_{5}}{4}+\frac{p_{6}}{2})(\frac{q_{11}}{4}+\frac{q_{12}}{2}+\frac{q_{8}}{4}+\frac{q_{9}}{2})v_{9}$ | [4] |
| *T* q_8_ _[t+1]_ = $((\frac{p_{2}}{2}+p_{3}+\frac{p_{5}}{4}+\frac{p_{6}}{2})(\frac{q_{10}}{2}+\frac{q_{11}}{4}+\frac{q_{7}}{2}+\frac{q_{8}}{4})+(p_{1}+\frac{p_{2}}{2}+\frac{p_{4}}{2}+\frac{p_{5}}{4})(\frac{q_{11}}{4}+\frac{q_{12}}{2}+\frac{q_{8}}{4}+\frac{q_{9}}{2}))v_{8}$ | [5] |
| *T* q_7_ _[t+1]_ = $(p_{1}+\frac{p_{2}}{2}+\frac{p_{4}}{2}+\frac{p_{5}}{4})(\frac{q_{10}}{2}+\frac{q_{11}}{4}+\frac{q_{7}}{2}+\frac{q_{8}}{4})v_{7}$ | [6] |
| *T* q_6_ _[t+1]_ = $(\frac{p_{5}}{4}+\frac{p_{6}}{2}+\frac{p_{8}}{2}+p_{9})(\frac{q_{2}}{4}+\frac{q_{3}}{2}+\frac{q_{5}}{4}+\frac{q_{6}}{2})v_{6}$ | [7] |
| *T* q_5_ _[t+1]_ = $((\frac{p_{5}}{4}+\frac{p_{6}}{2}+\frac{p_{8}}{2}+p_{9})(\frac{q_{1}}{2}+\frac{q_{2}}{4}+(\frac{1}{2}-d)q_{4}+\frac{q_{5}}{4})+(\frac{p_{4}}{2}+\frac{p_{5}}{4}+p_{7}+\frac{p_{8}}{2})(\frac{q_{2}}{4}+\frac{q_{3}}{2}+\frac{q_{5}}{4}+\frac{q_{6}}{2}))v_{5}$ | [8] |
| *T* q_4_ _[t+1]_ = $(\frac{p_{4}}{2}+\frac{p_{5}}{4}+p_{7}+\frac{p_{8}}{2})(\frac{q_{1}}{2}+\frac{q_{2}}{4}+(\frac{1}{2}-d)q_{4}+\frac{q_{5}}{4})v_{4}$ | [9] |
| *T* q_3_ _[t+1]_ = $(\frac{p_{2}}{2}+p_{3}+\frac{p_{5}}{4}+\frac{p_{6}}{2})(\frac{q_{2}}{4}+\frac{q_{3}}{2}+\frac{q_{5}}{4}+\frac{q_{6}}{2})v_{3}$ | [10] |
| *T* q_2_ _[t+1]_ = $((\frac{p_{2}}{2}+p_{3}+\frac{p_{5}}{4}+\frac{p_{6}}{2})(\frac{q_{1}}{2}+\frac{q_{2}}{4}+(\frac{1}{2}-d)q_{4}+\frac{q_{5}}{4})+(p_{1}+\frac{p_{2}}{2}+\frac{p_{4}}{2}+\frac{p_{5}}{4})(\frac{q_{2}}{4}+\frac{q_{3}}{2}+\frac{q_{5}}{4}+\frac{q_{6}}{2}))v_{2}$ | [11] |
| *T* q_1_ _[t+1]_ = $(p_{1}+\frac{p_{2}}{2}+\frac{p_{4}}{2}+\frac{p_{5}}{4})(\frac{q_{1}}{2}+\frac{q_{2}}{4}+(\frac{1}{2}-d)q_{4}+\frac{q_{5}}{4})v_{1}$ | [12] |
| *T* p_9_ _[t+1]_ = $(\frac{p_{5}}{4}+\frac{p_{6}}{2}+\frac{p_{8}}{2}+p_{9})(\frac{q_{11}}{4}+\frac{q_{12}}{2}+\frac{q_{5}}{4}+\frac{q_{6}}{2})u_{9}$ | [13] |
| *T* p_8_ _[t+1]_ = $((\frac{p_{5}}{4}+\frac{p_{6}}{2}+\frac{p_{8}}{2}+p_{9})(\frac{q_{10}}{2}+\frac{q_{11}}{4}+(\frac{1}{2}+d)q_{4}+\frac{q_{5}}{4})+(\frac{p_{4}}{2}+\frac{p_{5}}{4}+p_{7}+\frac{p_{8}}{2})(\frac{q_{11}}{4}+\frac{q_{12}}{2}+\frac{q_{5}}{4}+\frac{q_{6}}{2}))u_{8}$ | [14] |
| *T* p_7_ _[t+1]_ = $(\frac{p_{4}}{2}+\frac{p_{5}}{4}+p_{7}+\frac{p_{8}}{2})(\frac{q_{10}}{2}+\frac{q_{11}}{4}+(\frac{1}{2}+d)q_{4}+\frac{q_{5}}{4})u_{7}$ | [15] |
| *T* p_6_ _[t+1]_ = $((\frac{p_{2}}{2}+p_{3}+\frac{p_{5}}{4}+\frac{p_{6}}{2})(\frac{q_{11}}{4}+\frac{q_{12}}{2}+\frac{q_{5}}{4}+\frac{q_{6}}{2})+(\frac{p_{5}}{4}+\frac{p_{6}}{2}+\frac{p_{8}}{2}+p_{9})(\frac{q_{2}}{4}+\frac{q_{3}}{2}+\frac{q_{8}}{4}+\frac{q_{9}}{2}))u_{6}$ | [16] |
| *T* p_5_ _[t+1]_ = $((\frac{p_{2}}{2}+p_{3}+\frac{p_{5}}{4}+\frac{p_{6}}{2})(\frac{q_{10}}{2}+\frac{q_{11}}{4}+(\frac{1}{2}+d)q_{4}+\frac{q_{5}}{4})+(p_{1}+\frac{p_{2}}{2}+\frac{p_{4}}{2}+\frac{p_{5}}{4})(\frac{q_{11}}{4}+\frac{q_{12}}{2}+\frac{q_{5}}{4}+\frac{q_{6}}{2})+(\frac{p_{5}}{4}+\frac{p_{6}}{2}+\frac{p_{8}}{2}+p_{9})(\frac{q_{1}}{2}+\frac{q_{2}}{4}+\frac{q_{7}}{2}+\frac{q_{8}}{4})+(\frac{p_{4}}{2}+\frac{p_{5}}{4}+p_{7}+\frac{p_{8}}{2})(\frac{q_{2}}{4}+\frac{q_{3}}{2}+\frac{q_{8}}{4}+\frac{q_{9}}{2}))u_{5}$ | [17] |
| *T* p_4_ _[t+1]_ = $((p_{1}+\frac{p_{2}}{2}+\frac{p_{4}}{2}+\frac{p_{5}}{4})(\frac{q_{10}}{2}+\frac{q_{11}}{4}+(\frac{1}{2}+d)q_{4}+\frac{q_{5}}{4})+(\frac{p_{4}}{2}+\frac{p_{5}}{4}+p_{7}+\frac{p_{8}}{2})(\frac{q_{1}}{2}+\frac{q_{2}}{4}+\frac{q_{7}}{2}+\frac{q_{8}}{4}))u_{4}$ | [18] |
| *T* p_3_ _[t+1]_ = $(\frac{p_{2}}{2}+p_{3}+\frac{p_{5}}{4}+\frac{p_{6}}{2})(\frac{q_{2}}{4}+\frac{q_{3}}{2}+\frac{q_{8}}{4}+\frac{q_{9}}{2})u_{3}$ | [19] |
| *T* p_2_ _[t+1]_ = $((\frac{p_{2}}{2}+p_{3}+\frac{p_{5}}{4}+\frac{p_{6}}{2})(\frac{q_{1}}{2}+\frac{q_{2}}{4}+\frac{q_{7}}{2}+\frac{q_{8}}{4})+(p_{1}+\frac{p_{2}}{2}+\frac{p_{4}}{2}+\frac{p_{5}}{4})(\frac{q_{2}}{4}+\frac{q_{3}}{2}+\frac{q_{8}}{4}+\frac{q_{9}}{2}))u_{2}$ | [20] |
| *T* p_1_ _[t+1]_ = $(p_{1}+\frac{p_{2}}{2}+\frac{p_{4}}{2}+\frac{p_{5}}{4})(\frac{q_{1}}{2}+\frac{q_{2}}{4}+\frac{q_{7}}{2}+\frac{q_{8}}{4})u_{1}$ | [21] |
| *T* = $(p_{1}+\frac{p_{2}}{2}+\frac{p_{4}}{2}+\frac{p_{5}}{4})(\frac{q_{1}}{2}+\frac{q_{2}}{4}+\frac{q_{7}}{2}+\frac{q_{8}}{4})u_{1}+((\frac{p_{2}}{2}+p_{3}+\frac{p_{5}}{4}+\frac{p_{6}}{2})(\frac{q_{1}}{2}+\frac{q_{2}}{4}+\frac{q_{7}}{2}+\frac{q_{8}}{4})+(p_{1}+\frac{p_{2}}{2}+\frac{p_{4}}{2}+\frac{p_{5}}{4})(\frac{q_{2}}{4}+\frac{q_{3}}{2}+\frac{q_{8}}{4}+\frac{q_{9}}{2}))u_{2}+(\frac{p_{2}}{2}+p_{3}+\frac{p_{5}}{4}+\frac{p_{6}}{2})(\frac{q_{2}}{4}+\frac{q_{3}}{2}+\frac{q_{8}}{4}+\frac{q_{9}}{2})u_{3}+((p_{1}+\frac{p_{2}}{2}+\frac{p_{4}}{2}+\frac{p_{5}}{4})(\frac{q_{10}}{2}+\frac{q_{11}}{4}+(\frac{1}{2}+d)q_{4}+\frac{q_{5}}{4})+(\frac{p_{4}}{2}+\frac{p_{5}}{4}+p_{7}+\frac{p_{8}}{2})(\frac{q_{1}}{2}+\frac{q_{2}}{4}+\frac{q_{7}}{2}+\frac{q_{8}}{4}))u_{4}+((\frac{p_{2}}{2}+p_{3}+\frac{p_{5}}{4}+\frac{p_{6}}{2})(\frac{q_{10}}{2}+\frac{q_{11}}{4}+(\frac{1}{2}+d)q_{4}+\frac{q_{5}}{4})+(p_{1}+\frac{p_{2}}{2}+\frac{p_{4}}{2}+\frac{p_{5}}{4})(\frac{q_{11}}{4}+\frac{q_{12}}{2}+\frac{q_{5}}{4}+\frac{q_{6}}{2})+(\frac{p_{5}}{4}+\frac{p_{6}}{2}+\frac{p_{8}}{2}+p_{9})(\frac{q_{1}}{2}+\frac{q_{2}}{4}+\frac{q_{7}}{2}+\frac{q_{8}}{4})+(\frac{p_{4}}{2}+\frac{p_{5}}{4}+p_{7}+\frac{p_{8}}{2})(\frac{q_{2}}{4}+\frac{q_{3}}{2}+\frac{q_{8}}{4}+\frac{q_{9}}{2}))u_{5}+((\frac{p_{2}}{2}+p_{3}+\frac{p_{5}}{4}+\frac{p_{6}}{2})(\frac{q_{11}}{4}+\frac{q_{12}}{2}+\frac{q_{5}}{4}+\frac{q_{6}}{2})+(\frac{p_{5}}{4}+\frac{p_{6}}{2}+\frac{p_{8}}{2}+p_{9})(\frac{q_{2}}{4}+\frac{q_{3}}{2}+\frac{q_{8}}{4}+\frac{q_{9}}{2}))u_{6}+(\frac{p_{4}}{2}+\frac{p_{5}}{4}+p_{7}+\frac{p_{8}}{2})(\frac{q_{10}}{2}+\frac{q_{11}}{4}+(\frac{1}{2}+d)q_{4}+\frac{q_{5}}{4})u_{7}+((\frac{p_{5}}{4}+\frac{p_{6}}{2}+\frac{p_{8}}{2}+p_{9})(\frac{q_{10}}{2}+\frac{q_{11}}{4}+(\frac{1}{2}+d)q_{4}+\frac{q_{5}}{4})+(\frac{p_{4}}{2}+\frac{p_{5}}{4}+p_{7}+\frac{p_{8}}{2})(\frac{q_{11}}{4}+\frac{q_{12}}{2}+\frac{q_{5}}{4}+\frac{q_{6}}{2}))u_{8}+ (\frac{p_{5}}{4}+\frac{p_{6}}{2}+\frac{p_{8}}{2}+p_{9})(\frac{q_{11}}{4}+\frac{q_{12}}{2}+\frac{q_{5}}{4}+\frac{q_{6}}{2})u_{9}+(p_{1}+\frac{p_{2}}{2}+\frac{p_{4}}{2}+\frac{p_{5}}{4})(\frac{q_{1}}{2}+\frac{q_{2}}{4}+(\frac{1}{2}-d)q_{4}+\frac{q_{5}}{4})v_{1}+(\frac{p_{4}}{2}+\frac{p_{5}}{4}+p_{7}+\frac{p_{8}}{2})(\frac{q_{10}}{2}+\frac{q_{11}}{4}+\frac{q_{7}}{2}+\frac{q_{8}}{4})v_{10}+((\frac{p_{5}}{4}+\frac{p_{6}}{2}+\frac{p_{8}}{2}+p_{9})(\frac{q_{10}}{2}+\frac{q_{11}}{4}+\frac{q_{7}}{2}+\frac{q_{8}}{4})+(\frac{p_{4}}{2}+\frac{p_{5}}{4}+p_{7}+\frac{p_{8}}{2})(\frac{q_{11}}{4}+\frac{q_{12}}{2}+\frac{q_{8}}{4}+\frac{q_{9}}{2}))v_{11}+(\frac{p_{5}}{4}+\frac{p_{6}}{2}+\frac{p_{8}}{2}+p_{9})(\frac{q_{11}}{4}+\frac{q_{12}}{2}+\frac{q_{8}}{4}+\frac{q_{9}}{2})v_{12}+((\frac{p_{2}}{2}+p_{3}+\frac{p_{5}}{4}+\frac{p_{6}}{2})(\frac{q_{1}}{2}+\frac{q_{2}}{4}+(\frac{1}{2}-d)q_{4}+\frac{q_{5}}{4})+(p_{1}+\frac{p_{2}}{2}+\frac{p_{4}}{2}+\frac{p_{5}}{4})(\frac{q_{2}}{4}+\frac{q_{3}}{2}+\frac{q_{5}}{4}+\frac{q_{6}}{2}))v_{2}+(\frac{p_{2}}{2}+p_{3}+\frac{p_{5}}{4}+\frac{p_{6}}{2})(\frac{q_{2}}{4}+\frac{q_{3}}{2}+\frac{q_{5}}{4}+\frac{q_{6}}{2})v_{3}+(\frac{p_{4}}{2}+\frac{p_{5}}{4}+p_{7}+\frac{p_{8}}{2})(\frac{q_{1}}{2}+\frac{q_{2}}{4}+(\frac{1}{2}-d)q_{4}+\frac{q_{5}}{4})v_{4}+((\frac{p_{5}}{4}+\frac{p_{6}}{2}+\frac{p_{8}}{2}+p_{9})(\frac{q_{1}}{2}+\frac{q_{2}}{4}+(\frac{1}{2}-d)q_{4}+\frac{q_{5}}{4})+(\frac{p_{4}}{2}+\frac{p_{5}}{4}+p_{7}+\frac{p_{8}}{2})(\frac{q_{2}}{4}+\frac{q_{3}}{2}+\frac{q_{5}}{4}+\frac{q_{6}}{2}))v_{5}+(\frac{p_{5}}{4}+\frac{p_{6}}{2}+\frac{p_{8}}{2}+p_{9})(\frac{q_{2}}{4}+\frac{q_{3}}{2}+\frac{q_{5}}{4}+\frac{q_{6}}{2})v_{6}+(p_{1}+\frac{p_{2}}{2}+\frac{p_{4}}{2}+\frac{p_{5}}{4})(\frac{q_{10}}{2}+\frac{q_{11}}{4}+\frac{q_{7}}{2}+\frac{q_{8}}{4})v_{7}+((\frac{p_{2}}{2}+p_{3}+\frac{p_{5}}{4}+\frac{p_{6}}{2})(\frac{q_{10}}{2}+\frac{q_{11}}{4}+\frac{q_{7}}{2}+\frac{q_{8}}{4})+(p_{1}+\frac{p_{2}}{2}+\frac{p_{4}}{2}+\frac{p_{5}}{4})(\frac{q_{11}}{4}+\frac{q_{12}}{2}+\frac{q_{8}}{4}+\frac{q_{9}}{2}))v_{8}+ (\frac{p_{2}}{2}+p_{3}+\frac{p_{5}}{4}+\frac{p_{6}}{2})(\frac{q_{11}}{4}+\frac{q_{12}}{2}+\frac{q_{8}}{4}+\frac{q_{9}}{2})v_{9}$ | [22] |

Here, *T* is such that all the p_i_’s and q_i_’s add up to 1.

The frequencies of driving X and suppressors in males and females can be estimated at any time point using the associated genotypic frequencies.

| $X_{M}^{D}$ = $\frac{q_{4}+q_{5}+q_{6}+q_{10}+q_{11}+q_{12}}{\sum_{i=1}^{12} q_{i}}$ | [23] |
| --- | --- |
| $X_{F}^{D}$ = $\frac{p_{4}+p_{5}+p_{6}+2\left( p_{7}+p_{8}+p_{9} \right)}{\sum_{i=1}^{9} {2p}_{i}}$ | [24] |
| $A_{M}^{S}$ = $\frac{q_{2}+q_{5}+q_{8}+q_{11}+2(q_{3}+q_{6}+q_{9}+q_{12})}{\sum_{i=1}^{12} {2q}_{i}}$ | [25] |
| $A_{F}^{S}$ = $\frac{p_{2}+p_{5}+p_{8}+2(p_{3}+p_{6}+p_{9})}{\sum_{i=1}^{9} {2p}_{i}}$ | [26] |
| $Y^{S}$ = $\frac{\sum_{i=7}^{12} q_{i}}{\sum_{i=1}^{12} q_{i}}$ | [27] |

**Scenario A:**

We assume that our initial population is at equilibrium for X^D^ and test for the invasion of a Y-linked suppressor. To obtain solutions for the initial equilibrium frequencies in our model, we simplified our model by setting all genotypic frequencies linked to the autosomal suppressor to zero (p_2_, p_3_, p_5_, p_6_, p_8_, p_9_, q_2_, q_3_, q_5_, q_6_, q_8_, q_9_, q_11_, q_12_ = 0). We also set $s_{M}^{D}$=0 to simplify our model.

Now, to obtain the exact conditions for the invasion of the fixation equilibrium for X^D^, we set $X_{M}^{D}$=1, $X_{F}^{D}$=1, and $Y^{S}$=0. The limiting conditions thus obtained for the successful invasion of a Y-linked suppressor are –

| $s^{Y}$ < $2\times d$ | [28] |
| --- | --- |
| and, $s_{F}^{D}$ < $-\frac{d}{\left( h^{D}-2 \right)\left( d+1 \right)+1}$ | [29] |

A Y-linked suppressor could not invade the population if both conditions are not simultaneously satisfied. The condition presented in equation 28 has been previously calculated and presented in Hall, 2004. While equation 29 was not presented in Hall, 2004, it can be easily calculated not just from the reduced version of our model, but also from the equilibrium solutions presented in Hall, 2004. We further attempted to simplify the conditions by ignoring *h^D^* and were able to obtain a simple condition that could explain the invasion patterns of a Y-linked suppressor in our simulations. From our simulations, we found that a Y-linked suppressor could not invade when –

| $s^{Y}$ > $s_{F}^{D}$ ≥ $2\times d$ | [30] |
| --- | --- |

However, this condition was sufficient but not necessary to prevent the invasion of a Y-linked suppressor. There were other cases where this condition is not met, but the Y-linked suppressor still could not invade the population. It should be noted that the condition presented in equation 30 was derived purely from simulations.

**Supplementary Figures:**

**Figure S1: Equilibrium frequency of driving X before invasion of Y-linked suppressor** [$h^{A}$=0, $h^{D}$=0 (recessive costs), $s_{M}^{D}=$0, $s^{A}$=0.5, $s^{Y}$=0.1, 0.5, 0.9]. Equilibrium frequency of X^D^ in males before introduction of Y^S^ into the population. **A**. Autosomal suppressor is present in the initial population. **B**. Autosomal suppressor is absent in the initial population (Note that the white boxes on the top right are the spaces where X^D^ nearly fixes in the population in the absence of a suppressor due to strong drive with no fitness cost, hence the population becomes extinct). ‘*’ represents the parameter space representing where Y-linked suppressor can invade the population.

**Figure S2:** **Equilibrium frequency of the autosomal suppressor vs equilibrium frequency of driving X in males before the introduction of a Y-linked suppressor into the population**. The data in this graph represents a part of the parameter space [$h^{D}$=0, $h^{A}$=0, $s_{M}^{D}$=0, $s^{Y}$=0.1, $s^{A}$=0, 0.2, 0.5, 1], and shows whether a Y-linked suppressor invades or not.

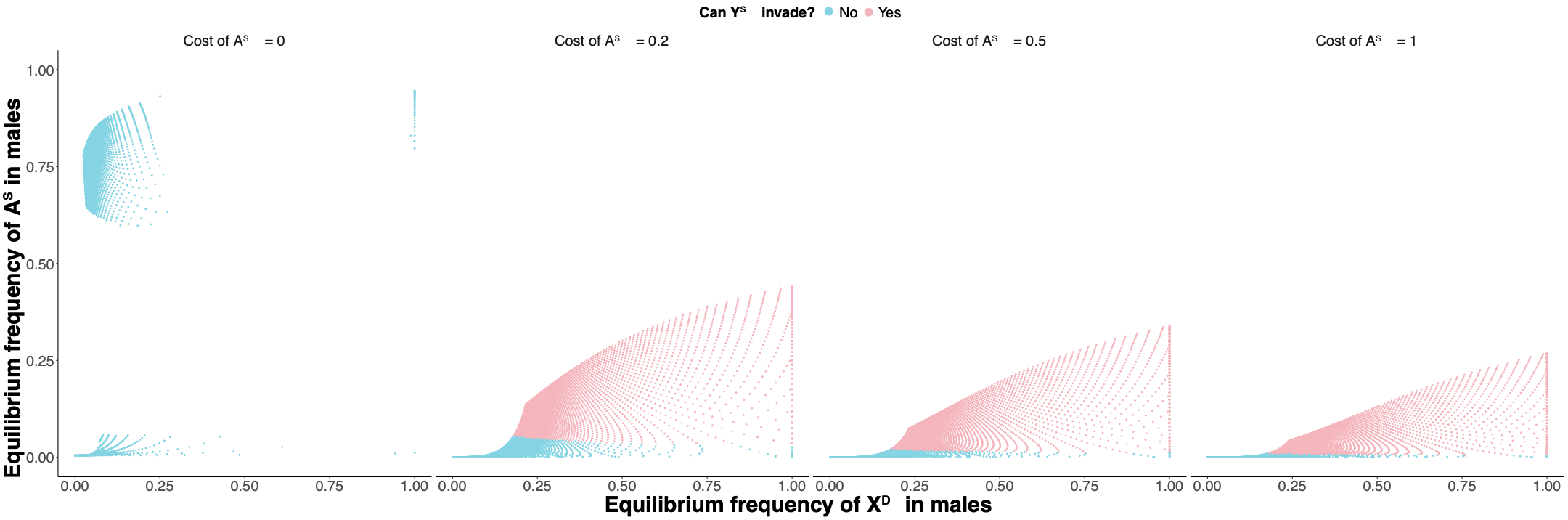

**Figure S3: Parameter space representing whether Y-linked suppressor can invade the population with variable dominance of the cost of driving X in females.** [$h^{A}$=0, $h^{D}$=0 (recessive), 0.5 (additive), 1 (dominant), $s_{M}^{D}$=0, $s^{A}$=0.5, $s^{Y}$=0.1].

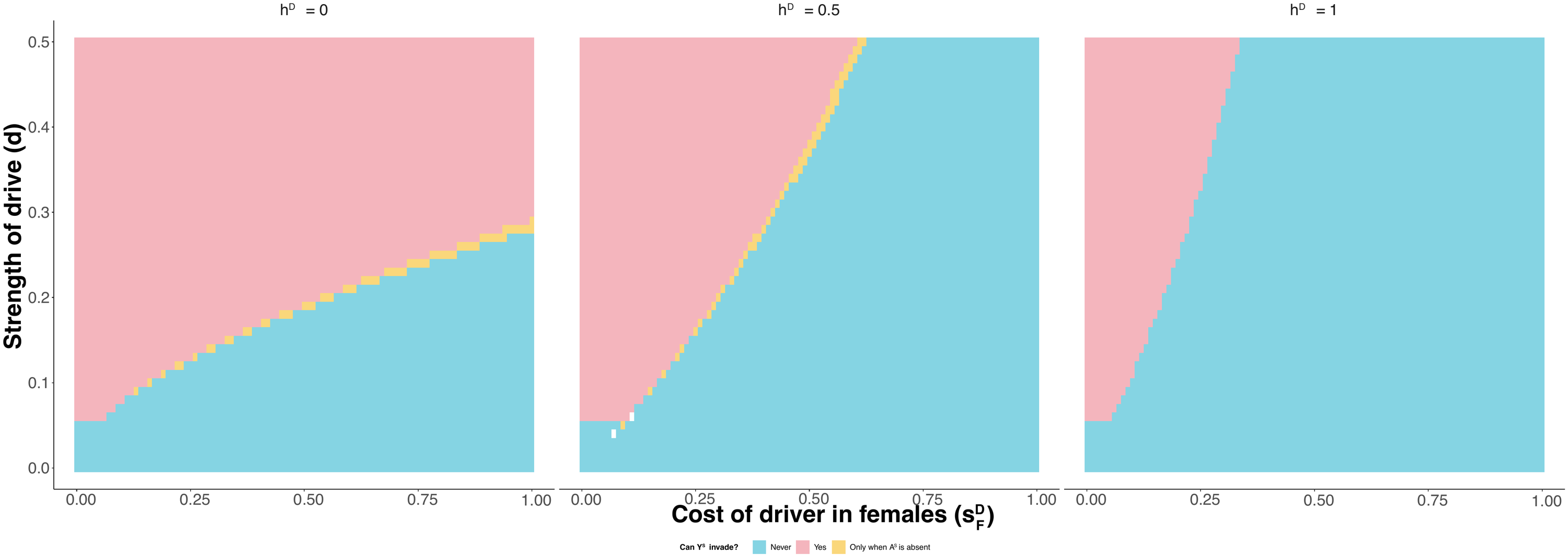

**Figure S4: Parameter space representing whether Y-linked suppressor can invade the population with variable dominance of the cost of suppression.** Autosomal suppressor is present in the initial population. [$h^{D}$=0, $h^{A}$=0 (recessive), 0.5 (additive), 1 (dominant), $s_{M}^{D}$=0, $s^{Y}$=0.5, $s^{A}$=0.1, 0.5, 0.9]

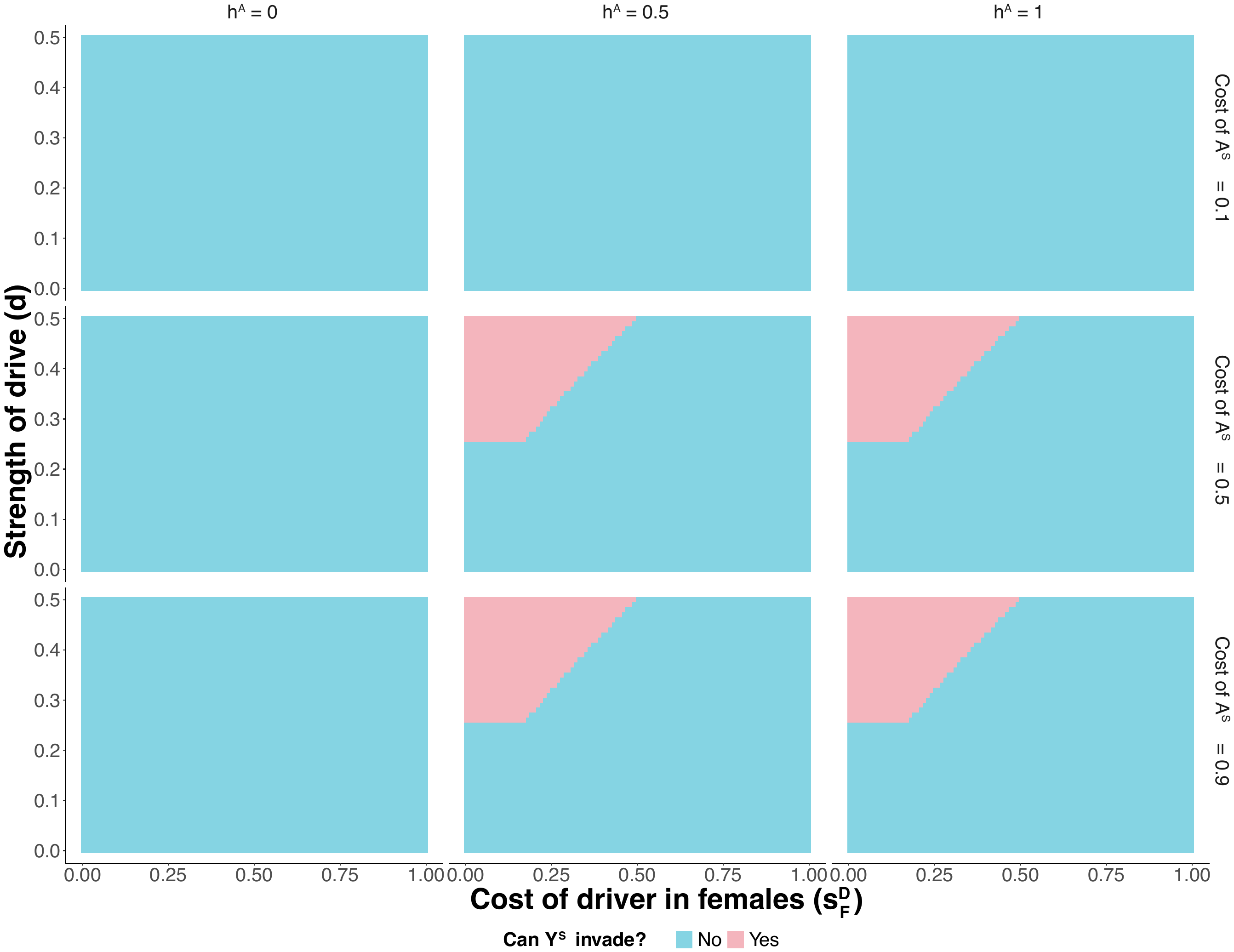

**Figure S5: Relative reduction in equilibrium frequencies of driving X upon invasion of a Y-linked suppressor** [$h^{A}$=0, $h^{D}$=0 (recessive costs), $s_{M}^{D}$=0, $s^{A}$=0.5, $s^{Y}$=0.1] **A**. Relative reduction in equilibrium frequency of X^D^ in males, **B.** Frequency of X^D^ in males before invasion of a Y-linked suppressor, **C.** Frequency of X^D^ in males after invasion of a Y-linked suppressor. The plots are faceted using the variable ‘YStable’ which implies whether the Y-linked suppressor was at equilibrium after invasion (YStable = Yes) or if it was cycling with the driving X (YStable = No).

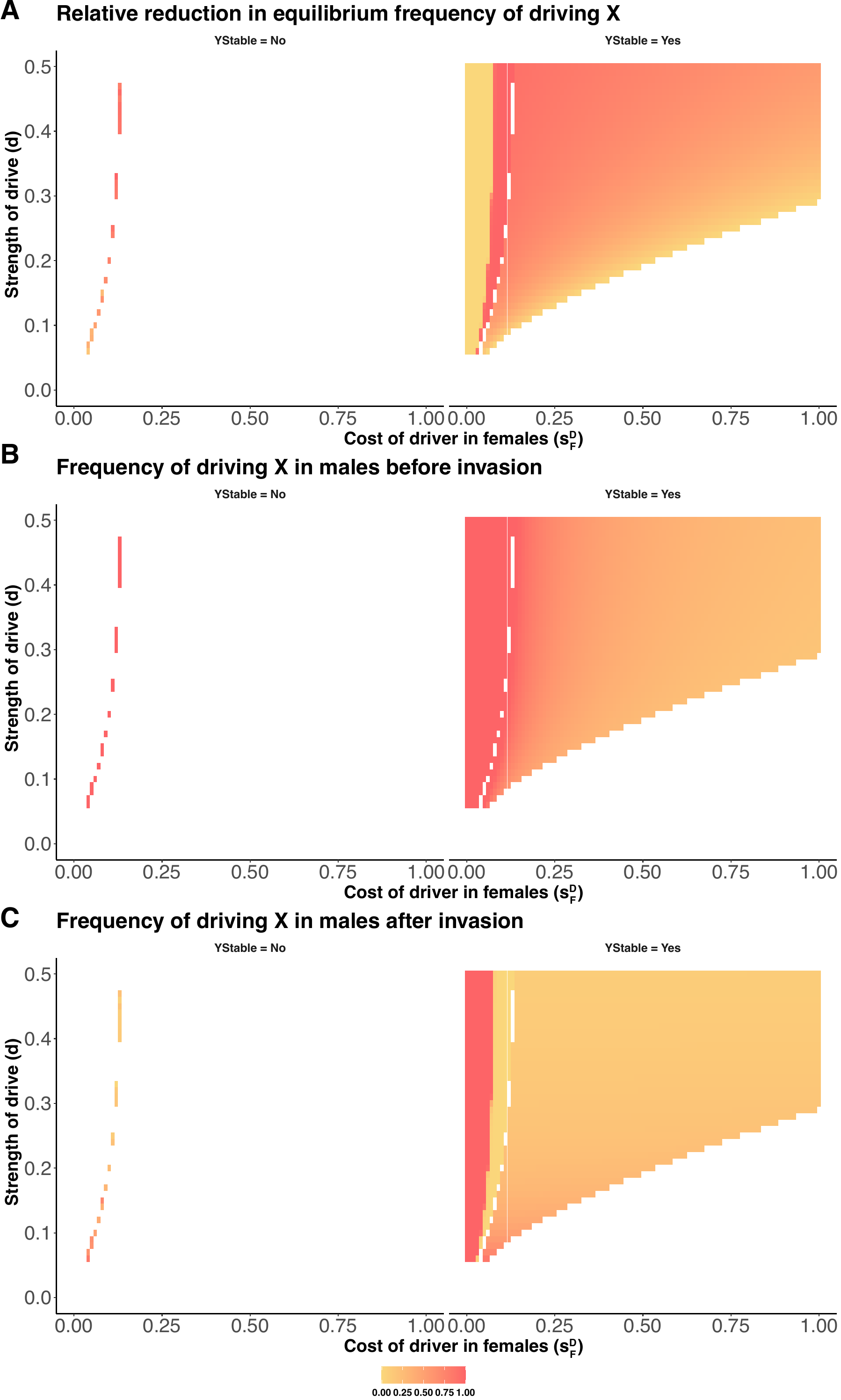

**Figure S6: Relative reduction in equilibrium frequencies upon invasion of a Y-linked suppressor** [$h^{A}$=0 (recessive), 0.5 (additive), 1 (dominant), $h^{D}$=0 (recessive costs), $s_{M}^{D}$=0, $s^{A}$=0.1, $s^{Y}$=0.1] **A**. Relative reduction in equilibrium frequency of X^D^ in males. **B**. Relative reduction in equilibrium frequency of A^S^ in males. The white space represents the space where Y^S^ cannot invade the population.

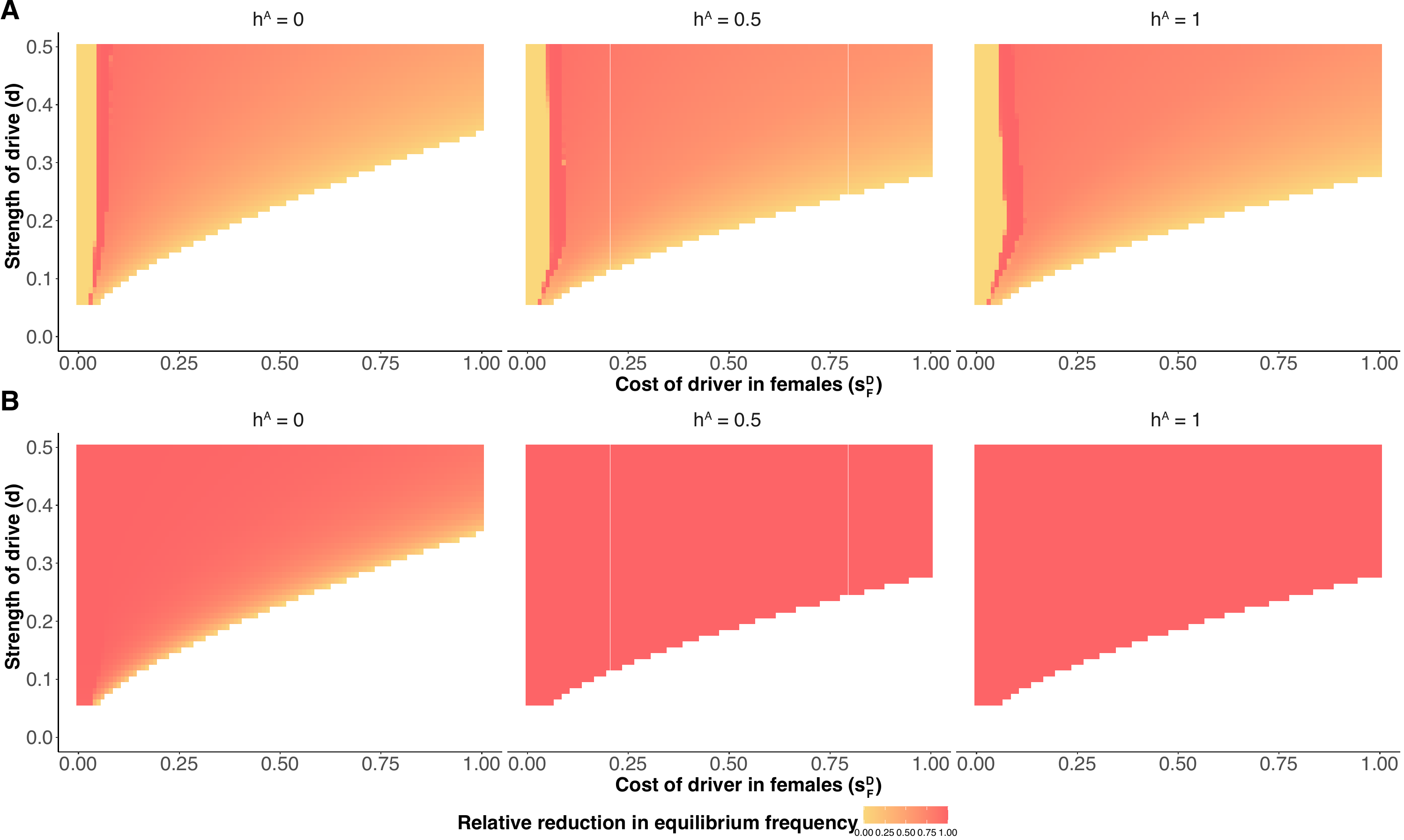

**Figure S7: Range of sex-ratios in the population at equilibrium for a Y-linked suppressor and driving X before invasion of an autosomal suppressor** [$h^{A}$=0, $h^{D}$=0 (recessive costs), $s_{M}^{D}$=0, $s^{A}$=0 (no cost), 0.5, 1, $s^{Y}$=0.1, 0.3, 1]. The ‘+’ sign indicates the parameter spaces where an autosomal suppressor could invade the population. A sex-ratio of <0.5 implies a male-biased sex ratio and a sex-ratio of >0.5 implies a female-biased sex-ratio. (Note that the white boxes corresponding to Figure 3 are the spaces where cycling occurs, and populations do not reach equilibrium as described by Hall 2004 and we deal with these parameter subsets separately in Scenario C).

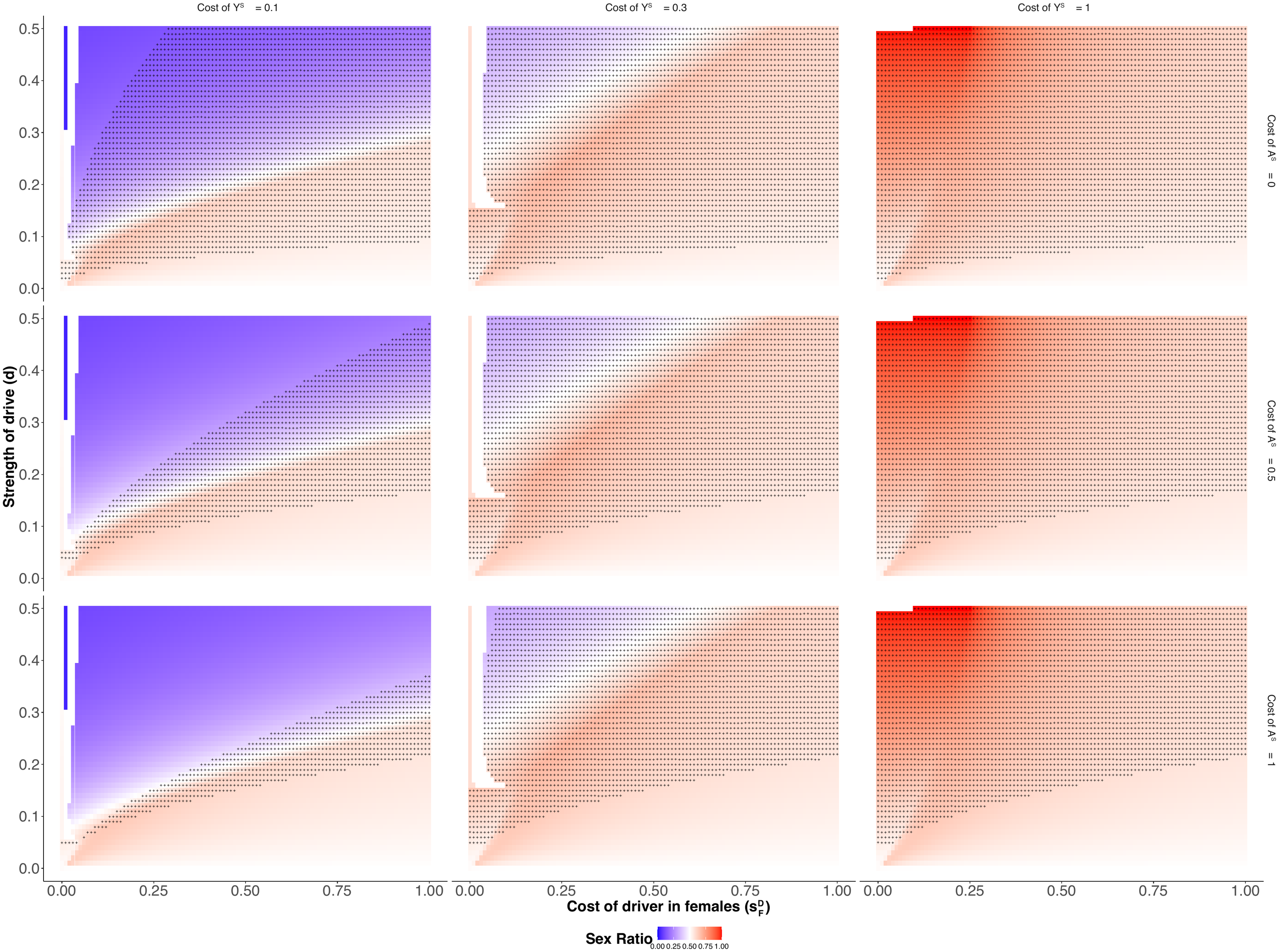

**Figure S8: Equilibrium frequency of a Y-linked suppressor (Y^S^) vs equilibrium frequency of driving X (X^D^) in males before introduction of autosomal suppressor (A^S^) into the population** [$h^{A}$=0, $h^{D}$=0 (recessive costs), $s_{M}^{D}$=0, $s^{A}$=0 (no cost), 0.5, 1, $s^{Y}$=0.1, 0.3, 1] The blue points represents the cases where an autosomal suppressor cannot invade into the population, and the pink points represents the cases where an autosomal suppressor can invade the population, for a range of costs of the autosomal and Y-linked suppressor (Y^S^).

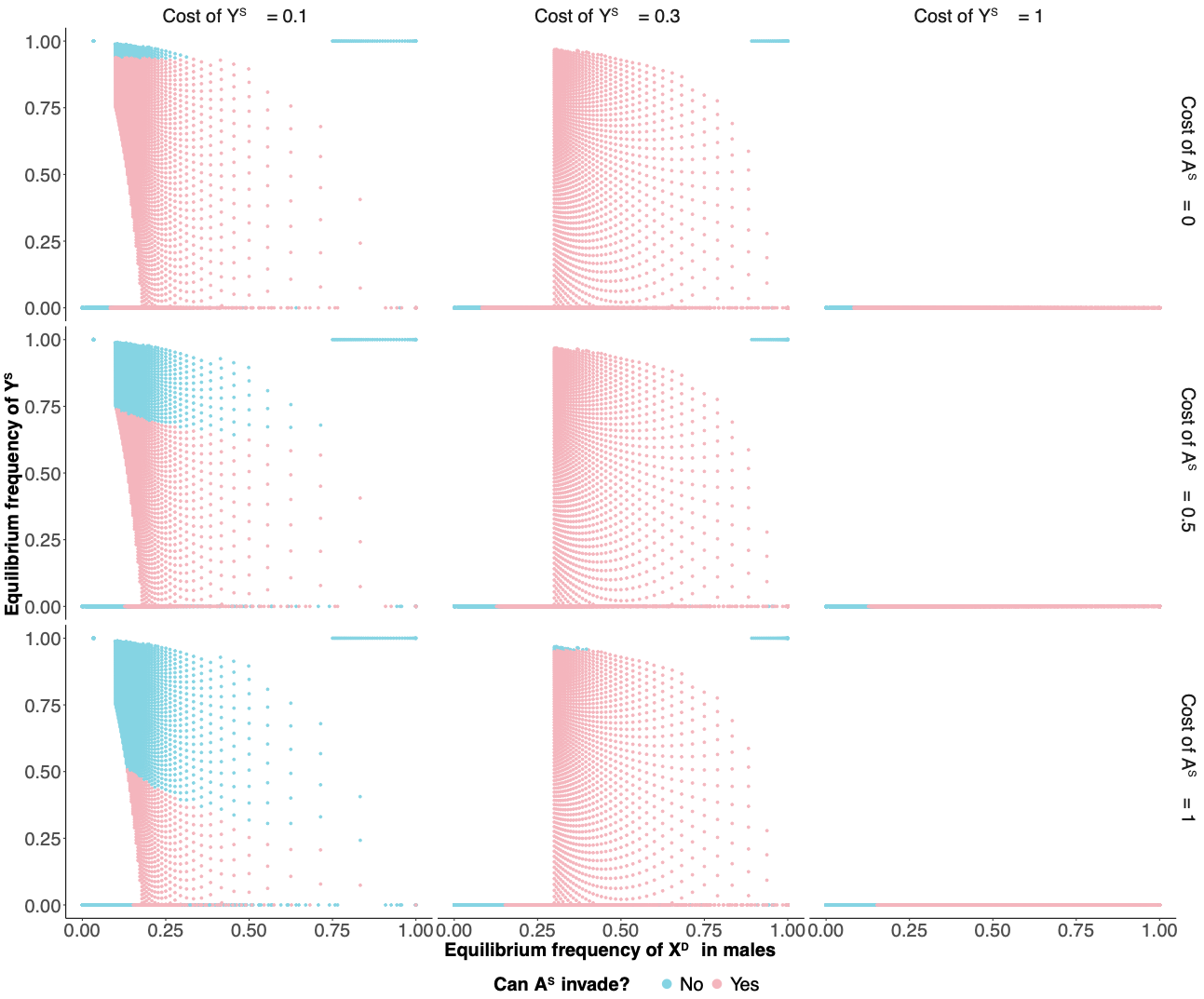

**Figure S9: Relative reduction in equilibrium frequencies upon invasion of an autosomal suppressor** [$h^{A}$=0, $h^{D}$=0 (recessive costs), $s_{M}^{D}$=0, $s^{A}$=0, 0.5, 1, $s^{Y}$=0.1, 0.3, 1] **A**. Relative reduction in equilibrium frequency of Y^S^ in males. **B**. Relative reduction in equilibrium frequency of X^D^ in males. The white space represents the space where A^S^ cannot invade the population. (Note that the white boxes corresponding to Figure 3 are the spaces where cycling occurs, and populations do not reach equilibrium as described by Hall 2004 and we deal with these parameter subsets separately in Scenario C).

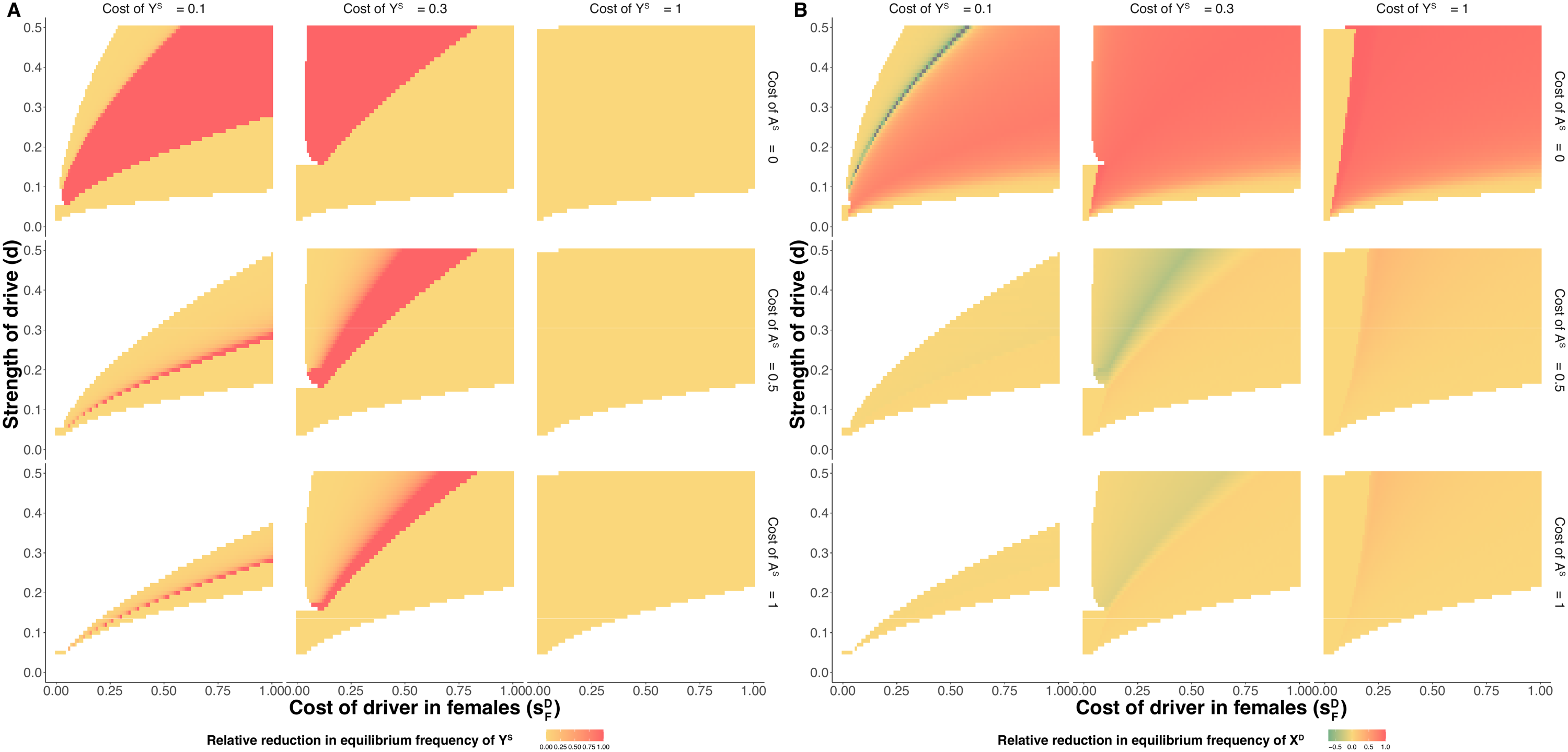

**Figure S10: Allelic frequencies vs time for populations in stable cycling for a Y-linked suppressor and driving X, with an equal sex-ratio before introduction of an autosomal suppressor** (measured at generation 5000). (Black: freq[Y^S^], blue: freq[X^D^], green: freq[A^S^]; solid lines: males, dashed lines: females; the red vertical line marks generation 5000, the timepoint of introduction of an autosomal suppressor into the population) **A.** $s_{M}^{D}$=0.2, $s_{F}^{D}$=0.1, $s^{A}$=0.3, $s^{Y}$=0.1, $h^{D}$=1, $h^{A}$=1, *d* =0.5; **B.** $s_{M}^{D}$=0.2, $s_{F}^{D}$=0.2, $s^{A}$=0.1, $s^{Y}$=0.1, $h^{D}$=1, $h^{A}$=1, *d* =0.5; **C.** $s_{M}^{D}$=0.3, $s_{F}^{D}$=0.1, $s^{A}$=0.1, $s^{Y}$=0.2, $h^{D}$=1, $h^{A}$=1, *d* =0.45; **D.** $s_{M}^{D}$=0.3, $s_{F}^{D}$=0.1, $s^{A}$=0.1, $s^{Y}$=0.3, $h^{D}$=1, $h^{A}$=1, *d* =0.4; **E.** $s_{M}^{D}$=0.3, $s_{F}^{D}$=0.1, $s^{A}$=0.2, $s^{Y}$=0.2, $h^{D}$=1, $h^{A}$=1, *d* =0.45; **F.** $s_{M}^{D}$=0.3, $s_{F}^{D}$=0.1, $s^{A}$=0, $s^{Y}$=0.3, $h^{D}$=1, $h^{A}$=1, *d* =0.4 (Note that for these separate subsets, we ran the simulations after introduction of an autosomal suppressor for an extra 20,000 generation to get a conclusive picture for the cycling dynamics and this figure represents only an example of some of these subsets).

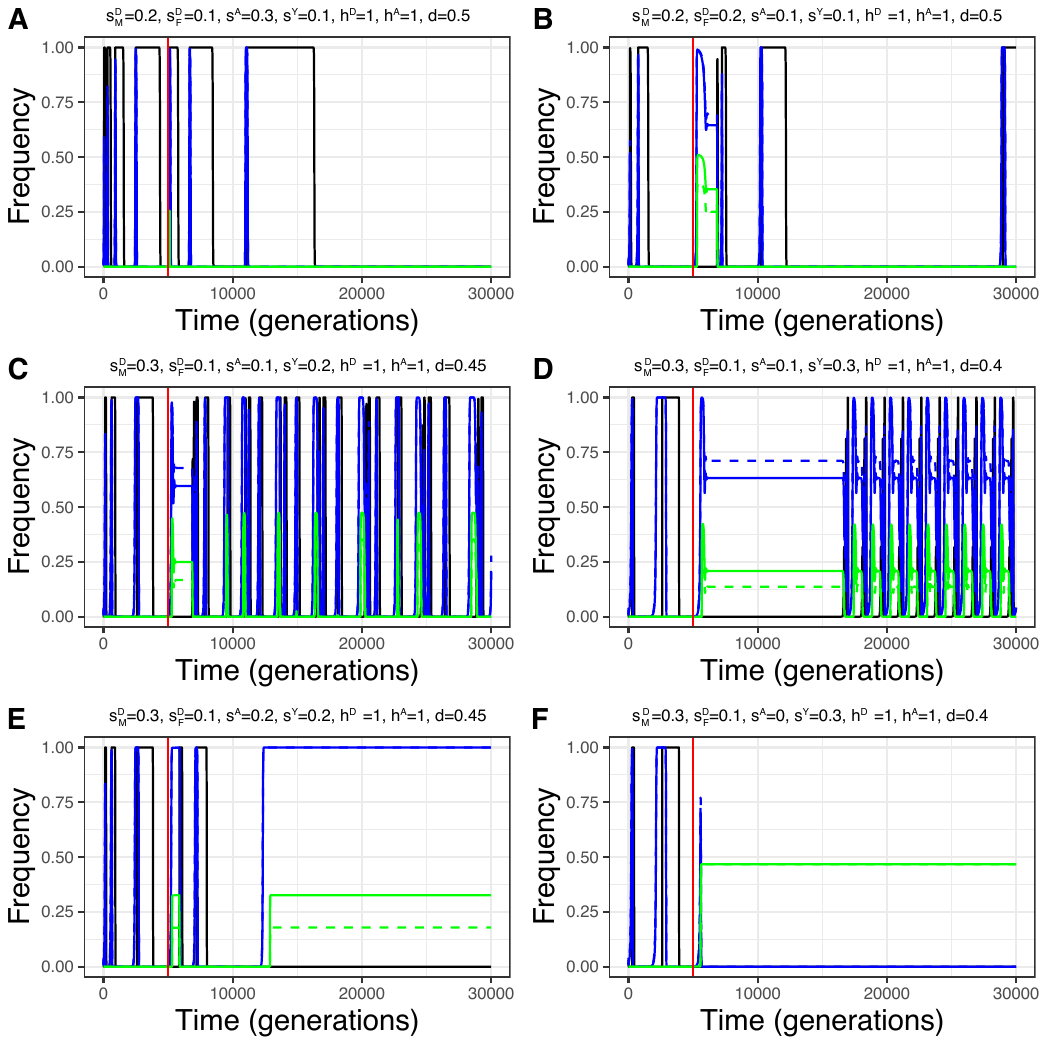

**Figure S11: Does cycling occur in populations with a finite population size?** Examples of regions of parameter spaces of stable cycling from Hall 2004, where we tested whether cycling persists in populations with a finite population size (10^6^). The cantaloupe color boxes represent the cases where cycling couldn’t occur in populations with a finite population size. The dark brown color boxes represent the cases where cycling persisted in population with a finite population size. (Note that these simulations were run for populations carrying a driving X and a Y-linked suppressor and were a separate set of simulations to include a finite population size in our infinite population size model)

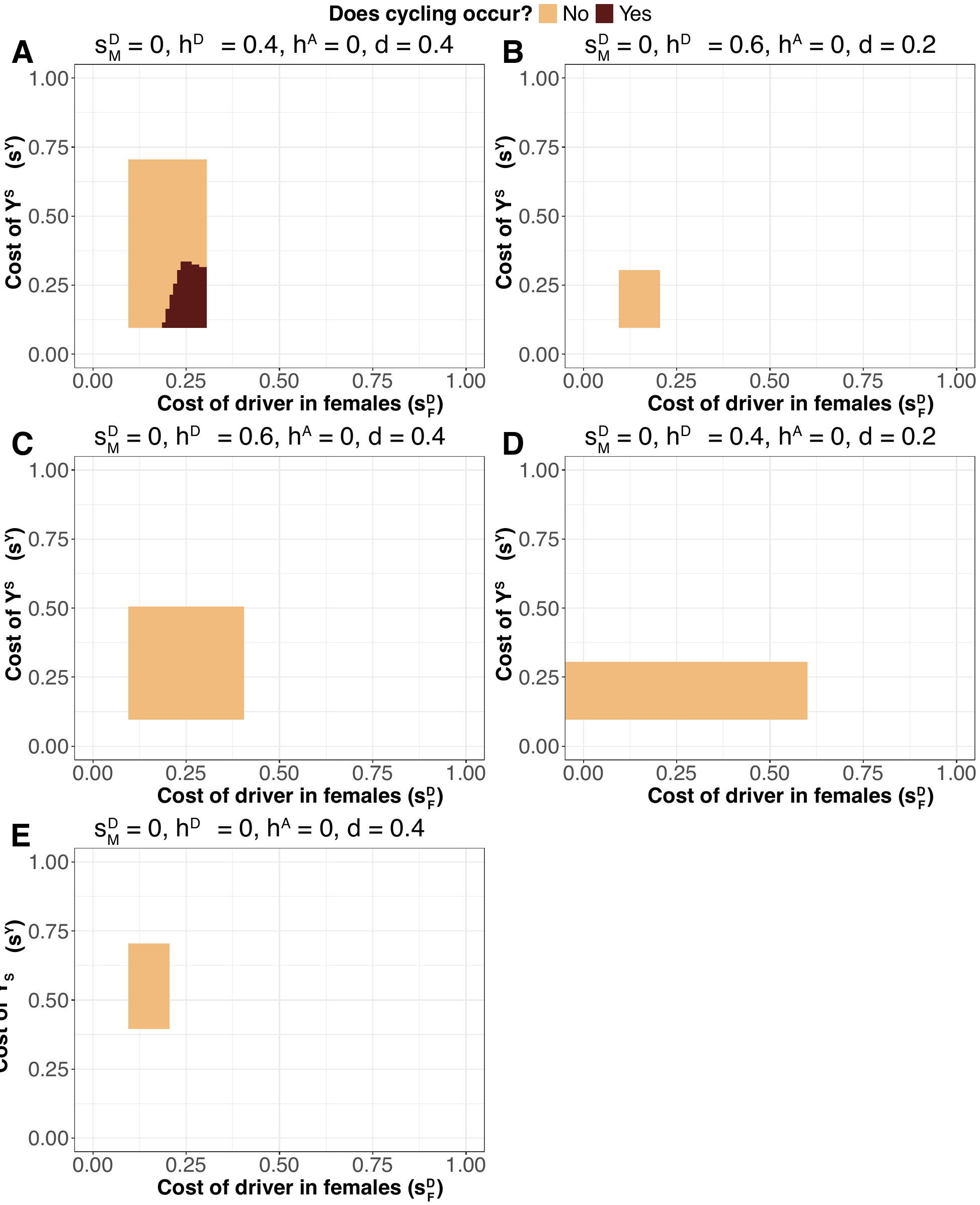
